# Differential regulation of BAX and BAK apoptotic activity revealed by a novel small molecule

**DOI:** 10.1101/2024.08.04.605933

**Authors:** Kaiming Li, Yu Q. Yap, Donia M. Moujalled, Fransisca Sumardy, Yelena Khakham, Angela Georgiou, Michelle Jahja, Thomas E. Lew, Melanie De Silva, Meng-Xiao Luo, Jia-nan Gong, Andrew W. Roberts, Zheng Yuan, Richard W. Birkinshaw, Peter E. Czabotar, Kym Lowes, David C. S. Huang, Benjamin T. Kile, Andrew H. Wei, Grant Dewson, Mark F. van Delft, Guillaume Lessene

**Affiliations:** The Walter and Eliza Hall Institute of Medical Research, Parkville, Victoria, Australia; Department of Medical Biology, The University of Melbourne, Parkville, Victoria, Australia; Department of Clinical Haematology, Peter MacCallum Cancer Centre and Royal Melbourne Hospital, Melbourne, Victoria, Australia; Laboratory of Human Disease Comparative Medicine, The Institute of Laboratory Animal Sciences, Chinese Academy of Medical Sciences & Peking Union Medical College; National Human Diseases Animal Model Resource Center; National Center of Technology Innovation for animal model, Beijing, China; Garvan Institute of Medical Research, Darlinghurst, Sydney, New South Wales, Australia; Department of Biochemistry and Pharmacology, The University of Melbourne, Parkville, Victoria, Australia

## Abstract

Defective apoptosis mediated by BAK or BAX underlies various human pathologies including autoimmune and degenerative conditions. The mitochondrial channel protein VDAC2 interacts with BAK and BAX through a common interface to either inhibit BAK or to facilitate BAX apoptotic activity. Using a newly developed small molecule (WEHI-3773) that inhibits the interaction between VDAC2 and BAK or BAX, we reveal contrasting effects on BAX and BAK apoptotic activity. WEHI-3773 inhibits apoptosis mediated by BAX by blocking VDAC2-mediated BAX recruitment to mitochondria. Conversely, WEHI-3773 primes BAK for apoptosis by impairing its inhibitory sequestration by VDAC2 on the mitochondrial membrane. In cells expressing both BAX and BAK, repressing their association with VDAC2 promotes apoptosis, because once BAK is activated, it further activates BAX through a feed-forward mechanism. In some leukemias, mutation or loss of BAX is a key driver of resistance to the BH3-mimetic anti-cancer drug venetoclax. Strikingly, promoting BAK-mediated killing by small molecule dissociation of the VDAC2 interaction can overcome this resistance in different leukemia models. These data reveal a hitherto unappreciated level of coordination of BAX and BAK apoptotic activity through their interaction with VDAC2 that may be targeted therapeutically.

## Introduction

Intrinsic or mitochondrial apoptosis is mainly governed by protein-protein interactions among BCL-2 proteins(*1*). To execute cell death, effector BCL-2 proteins BCL-2-associated X protein (BAX) or BCL-2 antagonist/killer (BAK) cause mitochondrial outer membrane permeabilization (MOMP), a critical event in the induction of apoptosis(*2*). Impaired mitochondrial apoptosis is a hallmark of cancer(*1*), while excessive cell death is observed in diverse conditions including ischemic brain injury and heart failure(*3, 4*). Targeting anti-apoptotic BCL-2 proteins to activate apoptosis has transformed the treatment of hematologic malignancies, as evidenced by the incorporation of venetoclax into standard of care for chronic lymphocytic and acute myeloid leukemia(*3*). BAK and BAX are pro-apoptotic effectors essential for the activity of venetoclax and other BH3-mimetic compounds. Venetoclax inhibits BCL-2 and thereby indirectly unleashes BAK and BAX to initiate apoptosis.

Venetoclax is a highly effective treatment for some blood cancers, but it is rarely curative and most patients relapse after an initial response(*5, 6*). Thus, acquired tolerance to venetoclax is a growing clinical problem and it is frequently associated with several adaptations that alter the expression or function of BCL-2 family members. Commonly observed changes include mutations to BCL-2 itself(*7*), elevated expression of the pro-survival proteins MCL-1 and/or BCL-X_L(*8*),_ as well as mutations that impair BAX function(*9*). Hence, there is a need for new drugs to target the apoptosis pathway at multiple levels and more effectively, either to enhance the killing effect of existing treatments like venetoclax, or to uncover new therapeutic vulnerabilities able to tackle the emerging problem of venetoclax resistance. One strategy of interest is to directly modulate the apoptosis effector proteins BAK and/or BAX and enhance their pro-apoptotic activity.

Alternatively, excessive apoptosis is observed in diverse conditions including ischemic brain injury, heart failure, and certain degenerative conditions(*3, 4*). It is unclear whether apoptosis inhibition will be a viable therapeutic strategy in these conditions(*3*) due to the lack of potent and specific small molecule inhibitors of BAK and/or BAX to test this hypothesis.

Directly targeting BAK and BAX to modulate apoptosis has proven challenging, partly due to the complex conformational and localization dynamics of these two proteins and their interactions with lipids within the mitochondrial outer membrane(*2*). Small molecules targeting BAK and BAX to modulate apoptosis have been reported mainly from target-based screening campaigns(*10-17*), with varying specificity, efficacy and mechanisms of action(*3, 16, 18-20*).

In addition to BCL-2 family proteins, several non-BCL-2 proteins interact with and regulate BAK and BAX. One well-established example is voltage dependent anion channel 2 (VDAC2)(*21*), which serves as an adapter to interact with BAK and BAX to regulate their mitochondrial localization and apoptotic activity(*22-26*). We report the discovery of a small molecule that blocks VDAC2 interactions with both BAX and BAK, leading to differential modulation of BAX and BAK apoptotic activity.

## Results

### High-throughput phenotypic screen identifies inhibitors of BAX-driven apoptosis

As published inhibitors of BAX generally lack specificity potency or defined mechanism of action, we undertook a cell-based phenotypic screen to identify inhibitors of BAX-mediated apoptosis. Our previous attempt at developing apoptosis inhibitors in cells expressing both BAX and BAK resulted in inhibitors of BAK-mediated apoptosis(*26*). While these compounds demonstrated potent and on-target activity, they were specific to mouse BAK and thus precluded application in human disease models. To identify inhibitors of BAX with selectivity and cellular potency in human cells, we engineered human myeloma cell line KMS-12-PE to be dependent on BAX (*BAK*^-/-^) or BAK (*BAX*^-/-^) by CRISPR/Cas9 editing (Fig. S1A). *BAK*^-/-^ or *BAX*^-/-^ KMS-12-PE cells remained sensitive to apoptotic cell death induced by the BH3-mimetic ABT-737 that inhibits the anti-apoptotic proteins BCL-2, BCL-X_L_ and BCL-W (Fig. S1B). Therefore, these cells provided suitable screening platforms to identify compounds able to inhibit either BAX- or BAK-mediated apoptosis.

High-throughput screening of our library comprising 120,652 compounds in *BAK*^-/-^ KMS-12-PE cells, followed by hit validation, led to identification of 17 distinct chemical series able to block BAX-mediated apoptosis induced by an EC_70_ dose of ABT-737 assessed by CellTiter-Glo^®^ (Fig. 1A). Detailed mechanistic studies concluded that 16 hit series failed to specifically modulate BAX or BAK dependent apoptosis. In particular, their lack of activity in the JC-1 mitochondrial depolarization assay suggested that their cell-based activity did not involve directly modulating the apoptotic machinery itself (*27*)(Fig. 1B and C). Only Compound **1** showed activity in both the primary cell viability assay using CellTiter-Glo^®^ and the mitochondrial depolarization assay (Fig. 1C-E). The inhibitory activity was specific to BAX as Compound **1** did not inhibit apoptosis in *BAX*^-/-^ or WT KMS-12-PE with functional BAK (Fig. S1C and S1D).

**Fig. 1.**
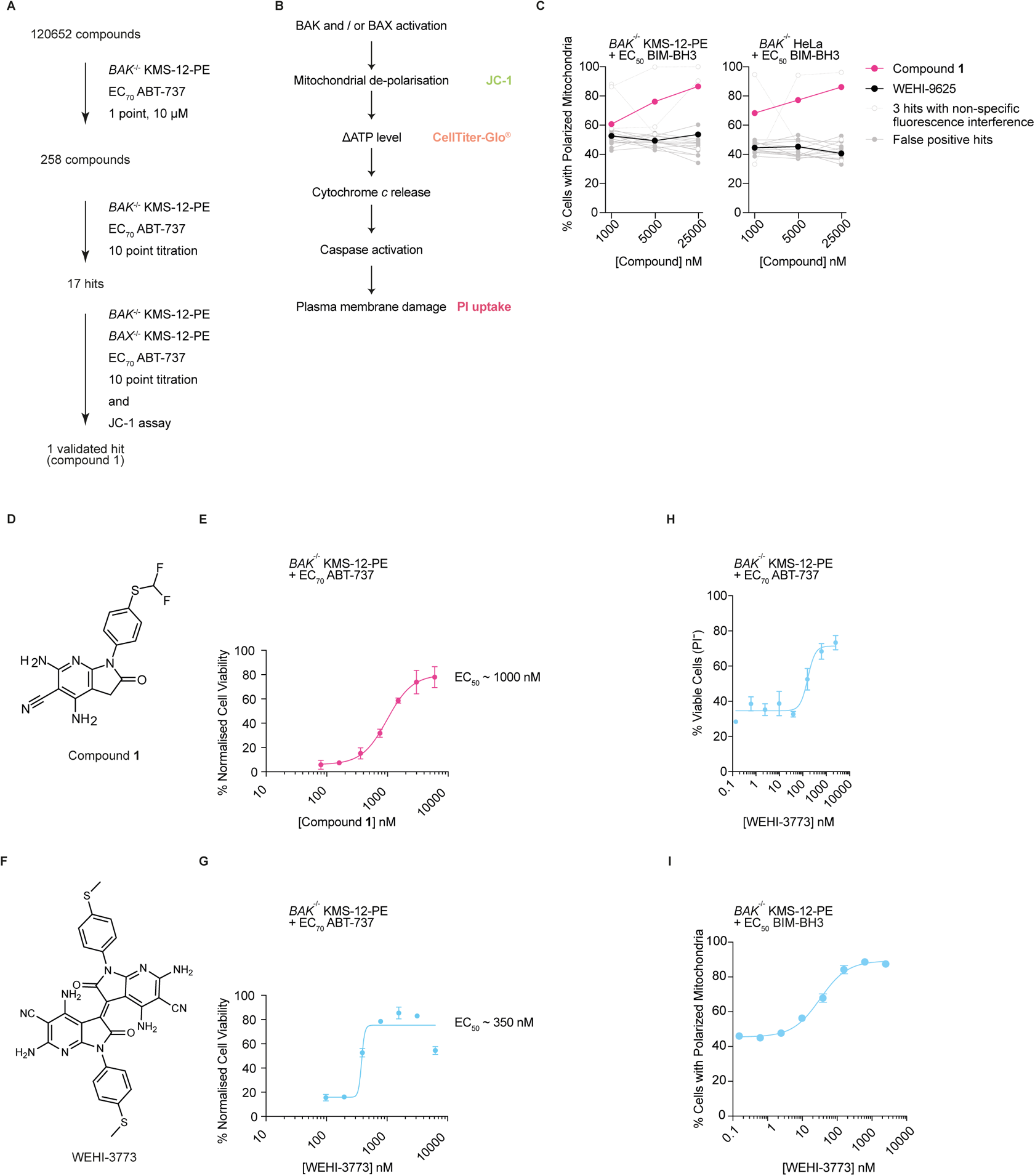
High-throughput screening identifies inhibitors of BAX-driven apoptosis. **(A)** Experimental workflow, and selection criteria employed for the discovery of inhibitors of human BAX-driven apoptosis. **(B)** Schematics of assays measuring different steps in the apoptosis pathway. **(C)** Dose-response curves for 17 hit compounds with mitochondria from *BAK*^-/-^ KMS-12-PE and *BAK*^-/-^ HeLa in JC-1 fluorescence assay. Data are from 1 independent experiment. **(D)** Chemical structures of Compound 1. **(E)** Dose-response curve for Compound 1 tested with *BAK*^-/-^ KMS-12-PE in the primary screen measured by CellTiter-Glo^®^. Data are mean ± S.E.M. of 2 independent experiments. **(F)** Chemical structures of WEHI-3773. **(G)** Dose-response curve for WEHI-3773 tested with *BAK*^-/-^ KMS-12-PE in the primary screen measured by CellTiter-Glo^®^. Data are mean ± S.E.M. of 2 independent experiments. **(H)** Dose response viability curves of *BAK*^-/-^ KMS-12-PE in the presence of EC_70_ concentration of ABT-737, measured by PI uptake and flow cytometry. Data are mean ± S.E.M. of 3 independent experiments. **(I)** Impact of WEHI-3773 on mitochondrial membrane potential measured by JC-1 staining and flow cytometry. Plasma-membrane-permeabilized *BAK*^-/-^ KMS-12-PE were co-treated with EC_50_ concentrations of human BIM-BH3 peptides. Data are mean ± S.E.M. of 3 independent experiments.

Intriguingly, the re-ordered Compound **1** sample developed increasing activity over time in our primary cell viability assay. Mass spectrometry indicated several species with molecular weights consistent with oxidation and/or dimerization (Fig. S1E). Using a similar substrate analogue to screening hit **1**, thiomethyl compound **2**, we were able to reproduce the decomposition profile observed in the screening sample (Fig. S1F) and isolate various components of this mixture. The minor fraction corresponding to a molecular weight of 619 (corresponding to 2M-4 of starting compound **2**) exhibited very high potency at blocking ABT-737-induced apoptosis in *BAK*^-/-^ KMS-12-PE cells with an EC_50_ less than 400 nM (Fig. 1F and G). Using structure determination by 2D NMR and other analytical methods, we determine that the active compound corresponded to compound WEHI-3773 (Fig. S1G, Supplemental Information). We found that a low-yielding, but reliable, method to obtain WEHI-3773 required dry DMSO and NaI (see Methods, Fig. S1G). All other protocols led to unworkable yields or various other isomers that were inactive in our primary assay.

As initial validation, our screening cascade (Fig. 1B) included orthogonal assays to measure cell viability by propidium iodide (PI) exclusion (Fig. 1H) or mitochondrial depolarization by JC-1 assay on enriched mitochondria (Fig. 1I). Cells were incubated with increasing concentration of WEHI-3773 together with BH3-mimetic compounds at a single EC_70_ concentration and viability or mitochondrial depolarization was assessed. In the viability assay, WEHI-3773 inhibited apoptosis in *BAK*^-/-^ KMS-12-PE cells with an EC_50_ comparable to that measured by CellTiter-Glo^®^ (Fig. 1H). In line with its increased cellular potency, WEHI-3773 also exhibited higher activity than Compound **1** in the JC-1 mitochondrial depolarization assay (Fig. 1I). These results confirmed the inhibitory effect of WEHI-3773 on BAX-driven apoptosis observed in the primary screening assay.

### WEHI-3773 inhibits BAX mitochondrial targeting and conformation change

Consistent with our observations in KMS-12-PE cells (Fig. 1H and 1I), WEHI-3773 also inhibited BAX-driven cell death and mitochondrial depolarization in BAK^-/-^ HeLa (Fig. 2A and 2B, S2A and S2B). The inhibition of mitochondrial depolarization in these two cell lines suggested that WEHI-3773 blocked BAX-driven cell death upstream of MOMP. In line with this, WEHI-3773 enabled long-term clonogenic survival of *BAK*^-/-^ HeLa and *BAK^-/-^* HCT116 cell lines treated with BH3 mimetics ABT-737 and MCL-1 inhibitor S63845, whereas caspase inhibition with QVD-OPh did not (Fig. 2C and 2D; Fig. S2C and S2D).

**Fig. 2.**
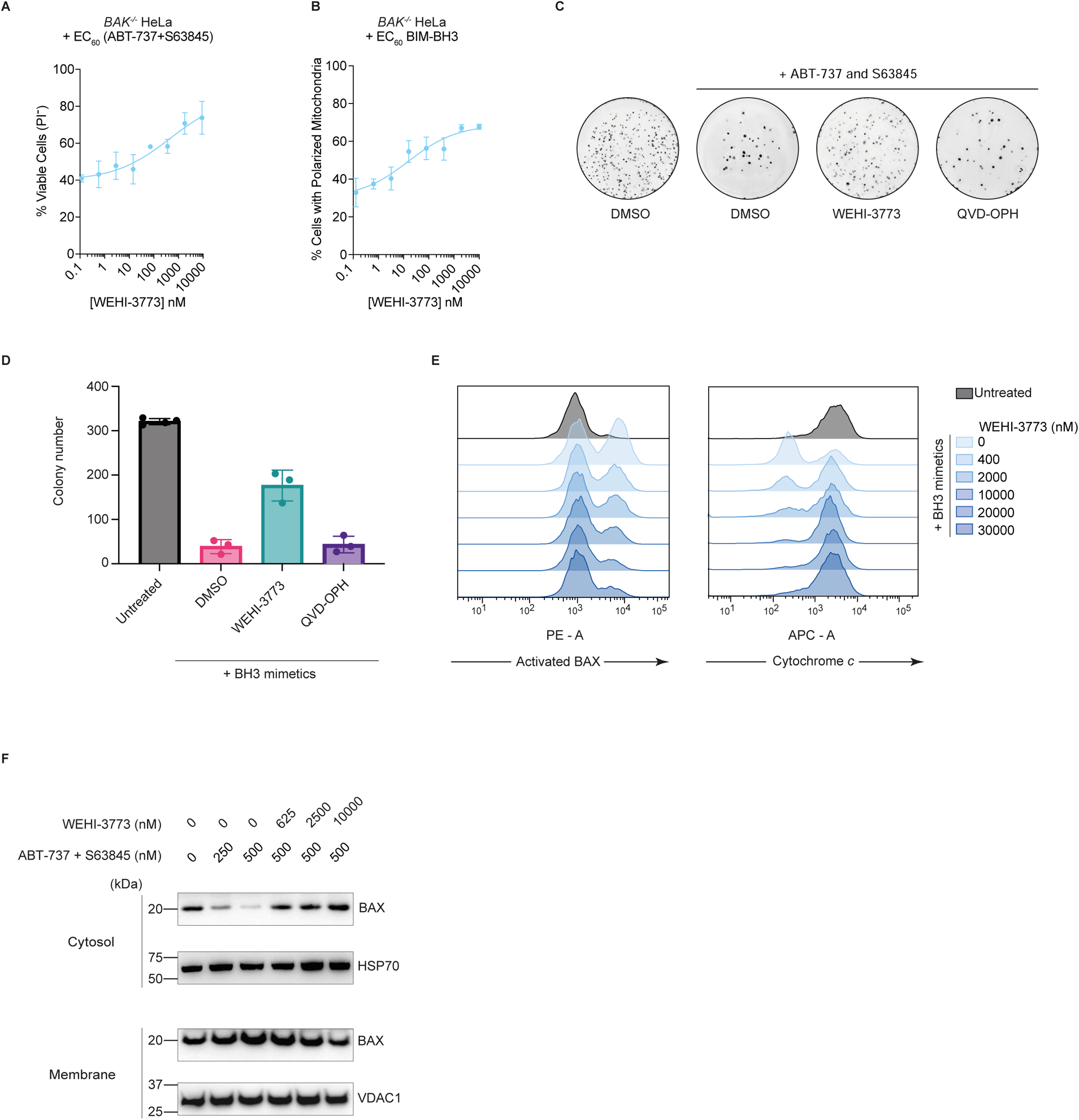
WEHI-3773 inhibits BAX activation and translocation to limit MOMP and protect cells. **(A)** Dose response viability curves of *BAK*^-/-^ HeLa in the presence of EC_60_ concentrations of ABT-737 and S63845, measured by PI uptake and flow cytometry. Data are mean ± S.E.M. of 3 independent experiments. **(B)** Impact of WEHI-3773 on mitochondrial membrane potential measured by JC-1 staining and flow cytometry. Plasma-membrane-permeabilized *BAK*^-/-^ HeLa were co-treated with EC_60_ concentrations of human BIM-BH3 peptides. Data are mean ± S.E.M. of 3 independent experiments. **(C)** Representative images of colony formation assay using *BAK*^-/-^ HeLa cells after 14 days of culture in the presence of ABT-737 and S63845 together with WEHI-3773, Q-VD.oph or DMSO. **(D)** Quantification of colony number. Data are as mean values and S.E.M of 3 independent experiments. **(E)** Intra-cellular staining, and flow cytometry to measure conformationally activated BAX and cytochrome *c* within *BAK*^-/-^ HeLa cells. **(F)** Western blot analysis of *BAK*^-/-^ HeLa cells treated with WEHI-3773, ABT-737 and S63845 at indicated concentrations for 2 hours. Representative blots from at least *n* = 3 independent experiments are shown.

During apoptosis, BAX undergoes structural changes to adopt an activated conformation that promotes MOMP and cytochrome *c* release. In doing so, BAX epitopes are exposed that can be recognized by conformation-specific antibodies(*28*). With these antibody tools, we observed that WEHI-3773 provided dose-dependent fashion inhibition of BAX conformational activation induced by combined BH3 mimetics, and this correlated with a blockade of mitochondrial cytochrome *c* release (Fig. 2E). Another critical step for BAX to mediate MOMP is its re-distribution from the cytosol to mitochondria. Notably, WEHI-3773 also blocked the redistribution of BAX to mitochondria during apoptosis induced by BH3 mimetics, indicating that WEHI-3773 interferes with this key early step in BAX-driven apoptosis (Fig. 2F).

### WEHI-3773 prevents VDAC2 from recruiting BAX to mitochondria

A proportion of BAX associates with mitochondria in healthy cells, even in the absence of an apoptotic stimulus. Notably, culturing cells with WEHI-3773 alone caused BAX to dissociate from mitochondria and be re-distributed to the cytosol (Fig. 3A). This phenomenon is reminiscent of the re-distribution of BAX in cells lacking VDAC2(*24, 29*) and led us to hypothesize that WEHI-3773 might interfere with the BAX:VDAC2 interaction and consequently the recruitment of BAX to mitochondria. Consistent with this hypothesis, WEHI-3773 was unable to provoke BAX re-distribution from mitochondria that lacked VDAC2 (Fig. 3A) and did not limit BAX-driven cell death in *Bak*^-/-^ *Vdac2*^-/-^ cells (Fig. 3B). To further assess the impact of WEHI-3773 on the BAX:VDAC2 interaction, we performed blue native polyacrylamide gel electrophoresis (BN-PAGE) on mitochondrial fractions isolated from *BAK^-/-^* HeLa cells treated with WEHI-3773. Strikingly, WEHI-3773 reduced the protein complex containing BAX and VDAC2 in a dose-dependent manner (Fig. 3C). Taken together, our data indicates that WEHI-3773 interferes with the BAX:VDAC2 interaction to inhibit BAX targeting to mitochondria (Fig. 3D).

**Fig. 3.**
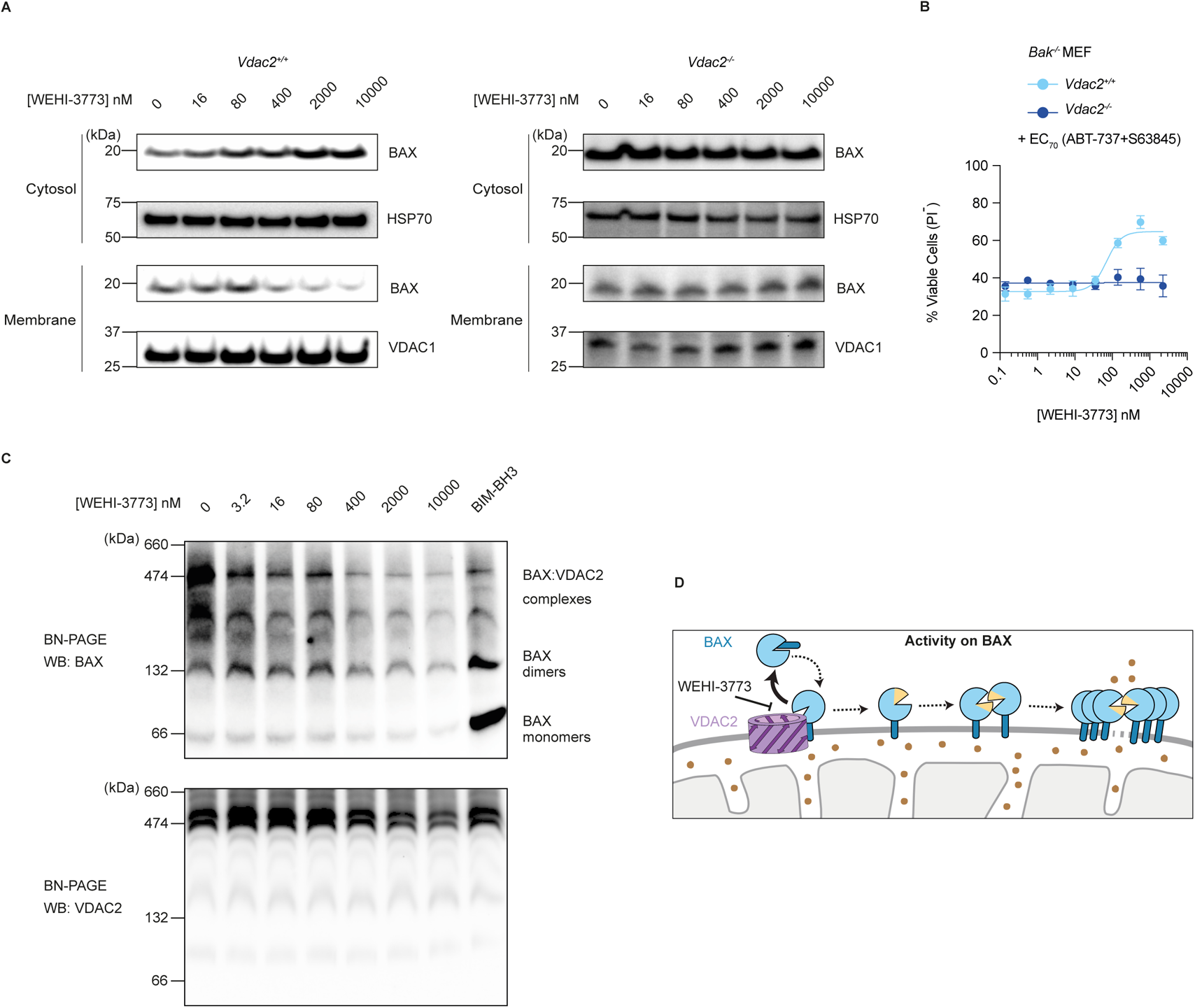
WEHI-3773 inhibits VDAC2-mediated BAX recruitment to mitochondria. **(A)** Western blot analysis of *Vdac2^-/-^* or *Vdac2^+/+^*MEF treated with WEHI-3773 alone at indicated concentrations for 2 hours. Representative blots of at least 3 independent experiments are shown. **(B)** Dose response viability curves of *Bak^-/-^* MEFs measured by PI uptake and flow cytometry. Cells were pre-treated with WEHI-3773 for 2 h first, then EC_70_ concentrations of ABT-737 plus S63845 were added for another 22 h before measurement. Data are mean ± S.E.M. of 3 independent experiments. **(C)** Western blot analysis of *BAK*^-/-^ mitochondria treated with WEHI-3773 or human BIM-BH3 peptide at indicated concentrations for 30 min, followed by blue native PAGE. Representative blots of at least 3 independent experiments are shown. **(D)** Proposed working model shows that WEHI-3773 acts on inhibiting BAX:VDAC2 interaction on mitochondria (gray) to limit MOMP and release of mitochondrial factors (brown dots).

### WEHI-3773 primes BAK activation by destabilizing its interaction with VDAC2

VDAC2 interacts with both BAX and BAK, but it regulates the apoptotic function of these proteins in opposing ways(*22, 24*). If WEHI-3773 inhibited BAX by interfering with the BAX:VDAC2 interaction, we hypothesized that it might also activate BAK if it similarly interfered with the BAK:VDAC2 interaction. To test this hypothesis, we evaluated the impact of Compound **1** and WEHI-3773 on BAK-driven apoptosis. Intriguingly, and consistent with our hypothesis, these compounds potentiated BAK-driven apoptosis in *BAX*^-/-^ and WT KMS-12-PE and HeLa cells (Fig. 4A and 4B). Furthermore, WEHI-3773 also potentiated BAK-driven, BIM-induced depolarization of mitochondria in *BAX*^-/-^ and WT HeLa (Fig. 4C). We confirmed that WEHI-3773 primed BAK conformational activation, as detected with conformation-specific antibody, and cytochrome *c* release in a dose-dependent manner (Fig. 4D). Furthermore, just as WEHI-3773 had the capacity to inhibit BAX in *Vdac2^+/+^*, but not *Vdac2^-/-^*MEF (Fig. 3A), it also primed BAK-mediated death in *Vdac2*^+/+^, but not *Vdac2*^-/-^ MEF (Fig. 4E). Thus, VDAC2 determines the capacity for WEHI-3773 to influence both BAX and BAK.

**Fig. 4.**
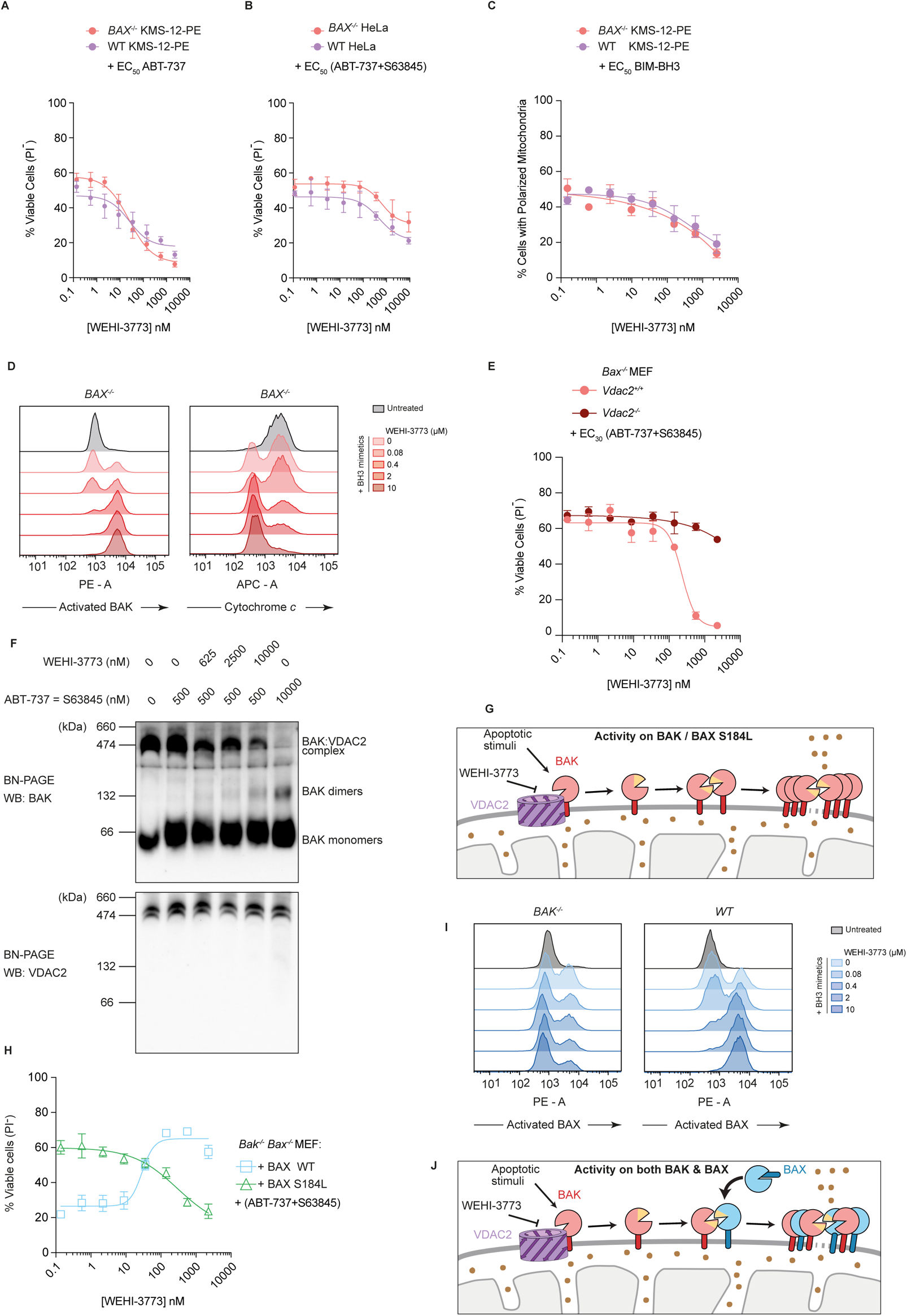
WEHI-3773 potentiates BAK activation by destabilizing the BAK:VDAC2 interaction. **(A)** Dose response viability curves of KMS-12-PE in the presence of EC_50_ concentration of ABT-737, measured by PI uptake and flow cytometry. Data are mean ± S.E.M. of 3 independent experiments. **(B)** Dose response viability curves of HeLa in the presence of EC_50_ concentrations of ABT-737 plus S63845, measured by PI uptake and flow cytometry. Data are mean ± S.E.M. of 3 independent experiments. **(C)** Impact of WEHI-3773 on mitochondrial membrane potential measured by JC-1 staining and flow cytometry. Plasma-membrane-permeabilized *BAX*^-/-^ or WT KMS-12-PE were co-treated with EC_50_ concentrations of human BIM-BH3 peptides. Data are mean ± S.E.M. of 3 independent experiments. **(D)** Dose response viability curves of *Bax^-/-^* MEF measured by PI uptake and flow cytometry. Cells were pre-treated with WEHI-3773 for 2 h, then EC_30_ concentrations of ABT-737 plus S63845 were added for another 22 h before measurement. Data are mean ± S.E.M. of 3 independent experiments. **(E)** Western blot analysis of *BAX^-/-^* HeLa treated with WEHI-3773, ABT-737 and S63845 at indicated concentrations for 2 h, followed by blue native PAGE. Representative blots of at least 3 independent experiments are shown. **(F)** Intra-cellular staining, and flow cytometry to measure conformationally activated BAK and preserved cytochrome *c* level within *BAX*^-/-^ HeLa cells. **(G)** Schematic model for WEHI-3773 inhibiting the interaction of VDAC2 with BAK / BAX S184L on mitochondria (gray) to potentiate MOMP and release of mitochondrial factors (brown dots) into cytosol. **(H)** Dose response viability curves of *Bak^-/-^ Bax^-/-^* MEFs re-expressing indicated BAX variants, measured by PI uptake and flow cytometry. Cells were pre-treated with WEHI-3773 for 2 h first, then EC_30_ or EC_70_ concentrations of ABT-737 plus S63845 were added for another 22 h before measurement. Data is mean ± S.E.M. of 3 independent experiments. **(I)** Intra-cellular staining and flow cytometry to measure conformationally activated BAX in *BAK*^-/-^ or WT HeLa. **(J)** Working model of WEHI-3773 in WT cells.

Notably, WEHI-3773 in the absence of BH3-mimetic treatment did not induce killing or activate BAK (Fig. S3A-C), suggesting that whilst it may not induce apoptosis in its own right, WEHI-3773 might synergize with, or potentiate, other sub-lethal stress signals. Consistent with these observations, WEHI-3773 alone did not dissociate BAK from VDAC2 (Fig. S3D), but it did prime BAK dissociation from VDAC2 when combined with low concentrations of BH3 mimetics (Fig. 4F and 4G).

To understand whether the differential effect of WEHI-3773 on BAX and BAK was due to their different sub-cellular localization, we tested the effect of the compound on BAX S184L, a mutant form of BAX that constitutively localizes to mitochondria(*28*). Like BAK, BAX S184L forms a stable complex with VDAC2 in healthy cells and dissociates from VDAC2 when activated (Fig. 4G)(*24*). As with its effect on BAK, but in contrast to its effect on wild-type BAX, WEHI-3773 potentiated apoptosis in *Bak*^-/-^ *Bax*^-/-^ MEF engineered to express BAX S184L (Fig. 4H), only when combined with sub-lethal apoptotic stress signals (Fig. S3E). Taken together, this data indicates that WEHI-3773 inhibits the interactions between VDAC2 and BAK or BAX (Fig. 4G). The distinct localization and pathway to activation on the mitochondrial outer membrane of these two proteins and their propensity to form stable interactions with VDAC2 may underpin the divergence in functional outcomes.

Interestingly, in cells expressing both BAK and BAX, BAX conformational activation was promoted, rather than inhibited by WEHI-3773 (Fig. 4I). This is likely due to the feed-forward amplification of BAX activation by activated BAK on mitochondria, a process that is independent of VDAC2 (Fig. 4J) (*30*). Given the fact that WEHI-3773 inhibits BAX in the absence of BAK, but promotes BAX activation when BAK is expressed or when BAX is targeted to mitochondria, we reasoned that WEHI-3773 did not interact with BAX or BAK directly to regulate their activity.

### WEHI-3773 targets VDAC2 to regulate BAK and BAX

Since WEHI-3773 could inhibit both BAK:VDAC2 and BAX:VDAC2 interactions, we investigated if WEHI-3773 directly engaged VDAC2 in cells. In our previous work, we discovered a small molecule, WEHI-9625, that interacts with VDAC2 to specifically inhibit mouse BAK. WEHI-9625 does not affect apoptosis driven by either human BAK or BAX(*26*). We have also shown using probes derived from WEHI-9625 that this compound directly interacts with both human and mouse VDAC2(*26*). We speculated that if WEHI-9625 and WEHI-3773 both interacted with VDAC2, they may do so competitively if their binding sites overlapped or were linked allosterically. If true, then although WEHI-9625 is not capable of influencing human BAK-driven or BAX-driven apoptosis on its own (Fig. S4A), it might interfere with the impact of WEHI-3773 on these proteins. Consistent with this, in KMS-12-PE engineered to lack either BAK or BAX, WEHI-9625 reduced the capacity of WEHI-3773 to both inhibit BAX-driven cell death and to prime BAK-driven apoptosis (Fig. 5A and B). That WEHI-9625 blocks WEHI-3773’s activity suggests that, like WEHI-9625, WEHI-3773 also engages VDAC2 in cells.

**Fig. 5.**
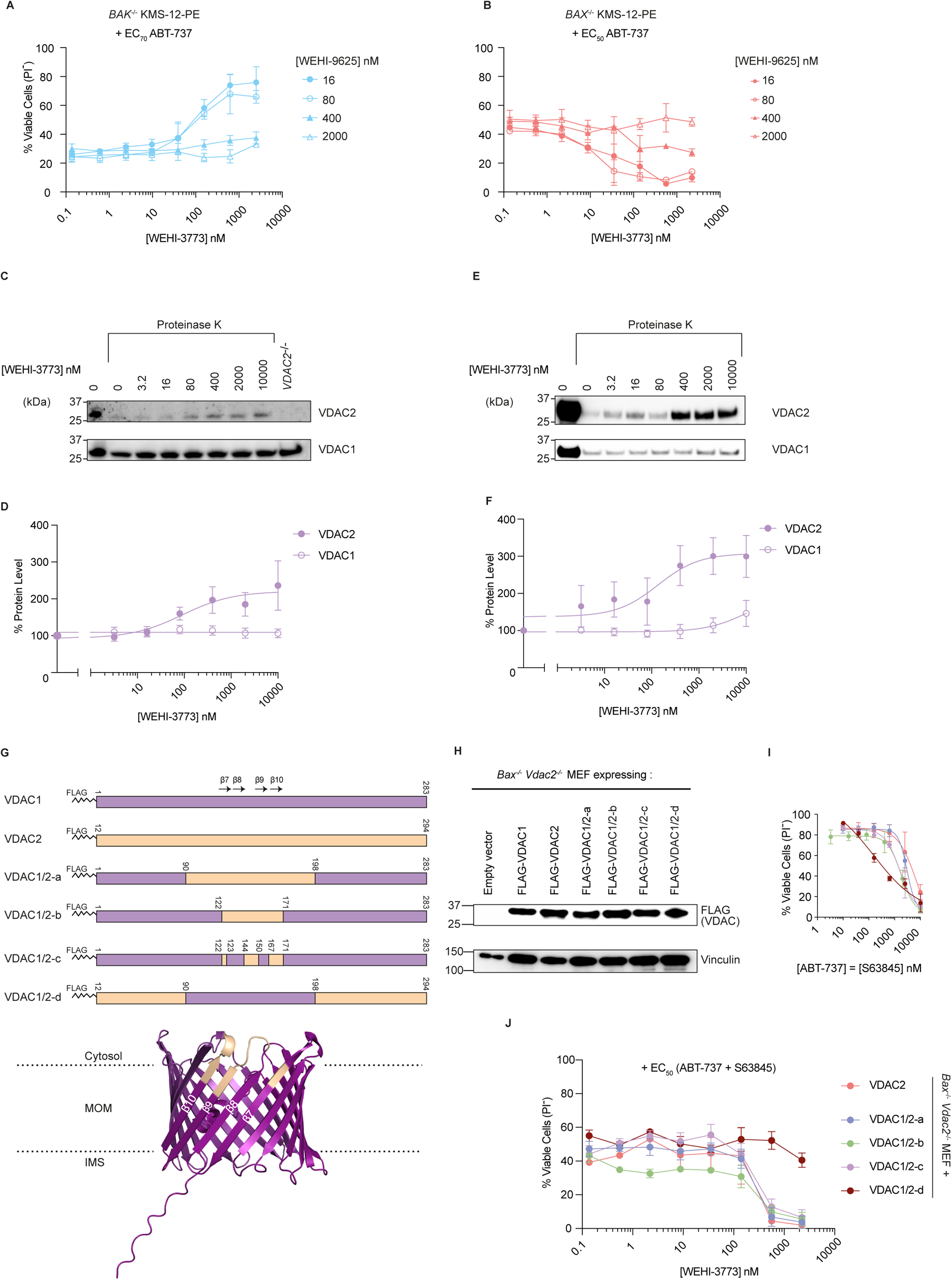
β7 to β10 of VDAC2 mediates the action of WEHI-3773. **(A)** Concentration-viability curves of *BAK*^-/-^ KMS-12-PE cells measured by PI uptake and flow cytometry. Cells were pre-treated with WEHI-3773 and WEHI-9625 for 2 h, then EC_70_ concentrations of ABT-737 was added for another 46 h before measurement. Data are mean ± S.E.M. of 3 independent experiments. **(B)** Dose response viability curves of *BAX*^-/-^ KMS-12-PE cells measured by PI uptake and flow cytometry. Cells were pre-treated with WEHI-3773 and WEHI-9625 for 2 h, then EC_50_ concentrations of ABT-737 was added for another 46 h before measurement. Data are shown as mean ± S.E.M. of 3 independent experiments. **(C)** Western blot analysis of mitochondria treated with WEHI-3773 at indicated concentrations for 30 min, followed by addition of proteinase K. Representative blots of 3 independent experiments are shown. **(D)** Immunoblots of (C) were quantified and normalized to the sample digested with proteinase K, but untreated with WEHI-3773. Data in the quantified plots are mean ± S.E.M. of 3 independent experiments. **(E)** Western blot analysis of recombinant proteins treated with WEHI-3773 or DMSO at indicated concentrations, followed by addition of proteinase K. Representative blots of 4 independent experiments are shown. **(F)** Immunoblots of (E) were quantified and normalized to the sample digested with proteinase K but untreated with WEHI-3773. Data in the quantified plots are mean ± S.E.M. of 4 independent experiments. **(G)** Schematic representation of VDAC1/VDAC2 chimeric constructs and cartoon presentation of the predicted human VDAC2 structure. VDAC2 is shown as cyan cartoons with the critical region examined in this study spanning β7 to β10 colored gold. **(H)** Western blot analysis of whole cell lysates of MEF cell lines. Representative blots of at least 3 independent experiments are shown. **(I)** Dose–response curves for ABT-737 plus S63845 on MEF re-expressing VDAC1 / VDAC2 chimeric proteins. Cells were treated with different concentrations of ABT-737 plus S63845. 24 h later, cells were collected, and cell death was quantified by PI uptake and flow cytometry. Data is mean ± S.E.M. of 3 independent experiments. **(J)** Dose response viability curves of *Bax^-/-^ Vdac2^-/-^* MEF re-expressing indicated VDAC variants, measured by PI uptake and flow cytometry. Cells were pre-treated with WEHI-3773 for 2 h, then EC_50_ concentrations of ABT-737 plus S63845 were added for another 22 h before measurement. Data is mean ± S.E.M. of 3 independent experiments.

To confirm the interaction between WEHI-3773 and VDAC2, we performed a drug affinity responsive target stability (DARTS) assay(*31*), based on the assumption that WEHI-3773 binding to VDAC2 would impact VDAC2 stability against proteolysis. Incubation of mitochondria isolated from HCT116 cells with WEHI-3773 inhibited proteolysis of VDAC2 in a dose-dependent manner, supporting the engagement of VDAC2 by WEHI-3773 on the mitochondrial membrane (Fig. 5C and 5D). Consistently, incubation of mitochondria with 400 nM WEHI-3773 inhibited proteolysis of VDAC2 with increasing concentrations of proteinase K (Fig. S4B and S4C).

We next sought to determine whether WEHI-3773 directly engages VDAC2 protein in solution or whether the native mitochondrial environment is required for the interaction. To this end, we expressed and purified recombinant full-length human VDAC1 and VDAC2 proteins from *E. coli.* (Fig. S5A and S5B). In line with the experiments in mitochondria, WEHI-3773 significantly inhibited proteolysis of recombinant VDAC2, but not VDAC1, (Fig. 5E and 5F). The above results suggest that WEHI-3773 targets VDAC2 both on mitochondria and in solution.

Based on the expression of VDAC1/VDAC2 chimeric proteins, previous studies identified β7 to β10 as the region of VDAC2 interacting with both BAK and BAX(*29, 32*) (Fig. 5G, Fig. S5C). This region on VDAC2 is also necessary for WEHI-9625 to inhibit mouse BAK-driven apoptosis(*26*). To further understand whether the β7 to β10 region on VDAC2 might mediate the action of WEHI-3773, we expressed VDAC1/VDAC2 chimeric proteins in *Bax*^-/-^*Vdac2*^-/-^ MEFs (Fig. 5H and 5I). Consistent with the competitive relationship between WEHI-9625 and WEHI-3773, substitution of VDAC1 β7 to β10 with the equivalent amino acid sequence from VDAC2 was sufficient to retain the capacity for WEHI-3773 to prime BAK-driven apoptosis (Fig. 5J). Conversely, replacement of that region in VDAC2 with the equivalent amino acid sequence from VDAC1 abolished its activity (Fig. 5J). Altogether, these results indicate that the region including β7 to β10 on VDAC2 is necessary and sufficient for the activity of WEHI-3773.

### WEHI-3773 overcomes venetoclax resistance in leukemic cells

Our previous studies identified loss-of-function BAX mutations as a mechanism of adaptive resistance to venetoclax-based therapy in patients with acute myeloid leukemia (AML)(*9*). Furthermore, we demonstrated that BAX-deficient leukemic cells were largely resistant to BCL-2-targeted compounds administered alone or in combination with other BH3 mimetics(*9*). We hypothesized that WEHI-3773 could re-sensitize BAX-deficient human leukemic cells to venetoclax by priming BAK activation. To examine this, we deleted BAX in MV4;11 cells by CRISPR/CAS9 gene editing (Fig. 6A) to render them resistant to venetoclax (Fig. 6B) and tested their responses to BH3 mimetics combined with WEHI-3773. At 200 nM, WEHI-3773 significantly sensitized these resistant MV4;11 cells to venetoclax (Fig. 6C), supporting our hypothesis that WEHI-3773 can overcome venetoclax resistance upon loss of BAX.

**Fig. 6.**
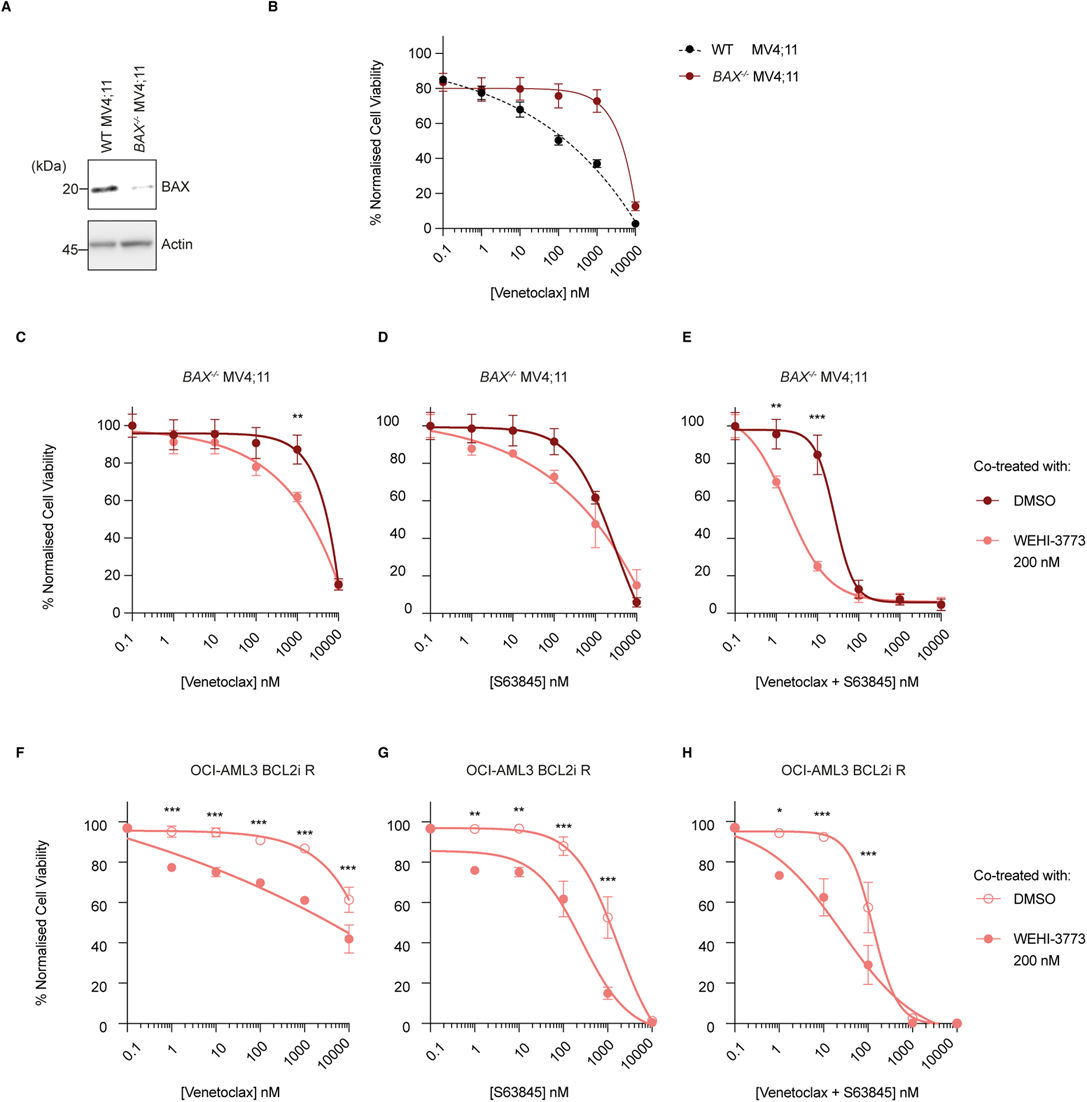
WEHI-3773 sensitizes BAX-mutant leukemia cells to BH3-mimetics. **(A)** Western blot of lysates from MV4;11 cells transduced with empty vector or *BAX* gRNA. **(B)** *BAX^-/-^* MV4;11 cells were treated with the indicated concentration of BH3 mimetics with or without WEHI-3773 (200 nM) prior to assessment of cell viability after 48 h. **(C-E)** *BAX^-/-^* MV4;11 cells were treated with the indicated concentration of venetoclax (C), S63845 (D) or combination of venetoclax and S63845 (E), with or without WEHI-3773 prior to assessment of cell viability after 48 h. Data is mean ± S.E.M. of 3 independent experiments. Two-way ANOVA with Tukey test; **p < 0.01, ***p < 0.001. **(F-H)** OCI-AML3 cells rendered resistant to venetoclax (*BCL-2i R*) were treated with the indicated concentration of venetoclax (F), S63845 (G) or combination of venetoclax and S63845 (H), with or without WEHI-3773 prior to assessment of cell viability after 48 h. Data is mean ± S.E.M. of 3 independent experiments. Two-way ANOVA with Tukey test; *p < 0.05, **p < 0.01, ***p < 0.001.

In addition to venetoclax, which has already been approved by FDA, MCL-1 inhibitors are also being tested in clinical trials for AML(*33*). In our previous study we also found that loss of BAX, but not its close relative BAK, could confer resistance to MCL-1 inhibitors in AML(*9*). WEHI-3773 could also sensitize resistant MV4;11 cells to MCL-1 inhibitor S63845 and the combination of venetoclax with S63845 (Fig. 6D and 6E), confirming that this compound can re-sensitize BAX-deficient human leukemic cells to inhibition of different anti-apoptotic BCL-2 proteins.

Given the promising effects of WEHI-3773 on BAX-deficient leukemic cells that are resistant to venetoclax, we then tested whether WEHI-3773 could promote apoptosis in a model of acquired resistance to BH3 mimetics(*9*). We have previously shown that OCI-AML3 leukemic cells exposed to prolonged treatment with BH3 mimetics acquired inactivating mutations in BAX that rendered them resistant to BCL-2 inhibitor (BCL-2i R), MCL-1 inhibitor and combination of both (BCL2i+MCL1i R) (Fig. S6A)(*9*). Hence, we tested whether WEHI-3773 could re-sensitize these resistant OCI-AML3 cells to BH3 mimetics. Indeed, WEHI-3773 significantly sensitized OCI-AML3 BCL-2i R cells to death induced by venetoclax (Fig. 6F), suggesting that WEHI-3773 could overcome acquired resistance to venetoclax. Additionally, WEHI-3773 could also sensitize OCI-AML3 BCL-2i R cells to MCL-1 inhibitor S63845 (Fig. 6G), or the combination of both (Fig. 6H). This data confirmed that WEHI-3773 can re-sensitize OCI-AML3 BCL-2i R cells to different BH3 mimetics. Likewise, WEHI-3773 sensitized OCI-AML3 cells rendered resistant to MCL-1 inhibition (MCL-1i R) to different BH3 mimetics (Fig. S6B). Overall, this data indicates that WEHI-3773 can restore the ability of BH3 mimetics, including venetoclax, to kill leukemic cells with acquired resistance.

The prolonged exposure of OCI-AML3 cells with combined BH3 mimetics (BCL2i + MCL1i) R selects for a loss-of-function BAK mutant(*9*). Thus, WEHI-3773 did not sensitize (BCL2i + MCL1i) R cells due to the BAK inactivation in these cells (Fig. S6C)(*9*). These results indicate that functional BAK capable of driving apoptosis is required for the action of WEHI-3773. In summary, our findings provide a proof-of-concept that small-molecule enhancement of BAK activity can overcome venetoclax resistance in different leukemic models.

## Discussion

Apoptosis mediated by BAK and BAX is aberrant in various human pathologies. Impaired mitochondrial apoptosis is a hallmark of cancer(*1*), while excessive cell death is observed in diverse conditions including ischemic brain injury and heart failure(*3, 4*). Hence, small molecules that can activate BAK- or BAX-driven apoptosis have the potential to improve treatments for cancer, while blocking these cell death effector proteins might rescue cells from degeneration.

Specific inhibitors of pro-survival BCL-2 proteins have entered the clinic(*34*). In contrast, the development of small molecules that specifically and directly target BAK or BAX to modulate apoptosis has proven more challenging. Several small molecule inhibitors of BAX and BAK have been reported, including recent report of covalent modifiers, that are providing new insight into the molecular control of cell death and the tractability of BAX and BAK to treat disease(*14-16, 20, 35*).

In this study, we used a cell-based phenotypic screen to identify a new class of small-molecules that differentially modulate both BAX and BAK through their shared interaction with VDAC2. A key feature of phenotypic drug discovery is the opportunity to identify small molecules with novel mechanisms of action. In the case of WEHI-3773, we identified a compound that both inhibits BAX and potentiates BAK-driven apoptosis. Mechanistically, WEHI-3773 targets VDAC2, a channel protein in the mitochondrial outer membrane that interacts with and regulates both BAK and BAX(*21, 24, 29*). Strikingly, its unique mechanism of action reveals the capacity to inhibit BAX only when BAK is expressed at low levels or the pro-apoptotic function of BAK is restricted. Notably, in certain cell types, such as differentiated neurons, deletion of BAX alone is sufficient to block apoptosis(*36*). WEHI-3773 would be a valuable tool in these systems to examine the cytoprotective effects of acute BAX inhibition.

As shown by the successful development of BH3 mimetics targeting pro-survival BCL-2 proteins, activation of apoptosis is a powerful strategy to treat some types of cancers(*37, 38*). By binding to pro-survival proteins, BH3 mimetics release the brakes on pro-apoptotic effectors BAK and BAX and thus activate them indirectly. Direct BAK or BAX activators might be another powerful anti-cancer strategy. However, molecules working in this way might induce broad toxicity. Since WEHI-3773 requires additional apoptotic stimuli to prime, but does not alone trigger BAK activity, the toxicity of drugs acting like WEHI-3773 may be manageable. As BAK and BAX are essential for the effective activity of BH3 mimetics, including venetoclax, small molecules that enhance their activity could synergize with BH3 mimetics and potentially overcome resistance to these compounds. By destabilizing the interaction between VDAC2 and BAK, WEHI-3773 is the first molecule to potentiate BAK activation through this mechanism. Hence, we envision that drug-like molecules with a similar mechanism of action to WEHI-3773 could be promising to address the issue of venetoclax resistance by enhancing BAK-dependent apoptosis.

In cells that express both apoptosis effectors, BAX and BAK, the net effect of WEHI-3773 is typically enhanced cell death, as the promotion of BAK-activity dominates the inhibitory effect on BAX. Moreover, since activated BAK can induce BAX integration into the mitochondrial outer membrane and conformational activation independently of VDAC2(*30*), WEHI-3773 effectively promotes both effectors to kill. However, this will likely be context dependent, because in cells with low or functionally impaired BAK, the BAX-inhibitory activity of WEHI-3773 may dominate.

Finally, our discovery of another VDAC2-targeting compound through a phenotypic drug screen further emphasized the pivotal role of VDAC2 in regulating BAK- and BAX-mediated apoptosis. In line with previous studies(*24, 26, 29*), this work shows that the same region on VDAC2 is responsible for engaging BAK and BAX, and that this region on VDAC2 can be targeted by small molecules. Interestingly, the fact that WEHI-3773 exhibits a mechanism that is distinct from our previously described BAK inhibitor WEHI-9625(*26*), while engaging a similar region of VDAC2, suggests that subtle differences in VDAC2 engagement lead to strikingly different effects on BAX’s and BAK’s ability to drive apoptosis. Moreover, we demonstrate that although BAK and BAX are challenging targets, modulating their interaction with VDAC2 might be an alternative strategy to affect their apoptotic activity. To this end, defining the molecular and structural details of the BAK:VDAC2 and BAX:VDAC2 interfaces will be informative(*21, 39*). We envision that research tools like WEHI-3773 will help us to uncover details of this interaction and move closer to the development of apoptosis-modulating therapeutics.

## Materials and Methods

### Constructs, antibodies, and reagents

The following constructs were used in this study: FLAG-tagged wild-type human BAX or FLAG-tagged human BAX S184L were cloned into pMSCV-IRES-GFP. pCMV-VSV-G (Addgene, 8454) and gag/pol (Addgene, 14887) were used for the generation of lentivirus for stable expression.

The following primary antibodies were used. BAK (B5897, Sigma-Aldrich), BAX (49F9, in-house from DCSH), VDAC2 (ab37985, Abcam), VDAC1 (ab37985, Millipore), HSP70 (MA3-006, Thermo Fisher Scientific) and cytochrome *c* (Clone#7H8.2C12, BD Pharmingen). Secondary antibodies anti-rabbit IgG (Cat#403005), anti-mouse IgG (Cat#103005), anti-rat IgG (Cat#303005) were obtained from Southern Biotech. BH3 mimetics venetoclax (cat# S8048, Selleckchem/Jomar Life Research), ABT-737 (Cat#A-1002, Active Biochem), A-1331852 (WEHI Chemical Biology Division) and S63845 (Cat#A-6044, Active Biochem) were used.

### Cell culture and transient transfection

SV40-transformed Mouse Embryonic Fibroblasts (MEFs) were cultured in Dulbecco’s Modified Eagle’s Medium (DMEM) supplemented with 10% (v/v) Fetal Bovine Serum (FBS), 50 μM 2-Mercaptoethanol (2-ME) and 100 μM asparagine. HeLa cells (ATCC #CCL-2) and Human Embryonic Kidney 293 T (HEK293T) cells (ATCC #CRK-3216) were cultured in DMEM supplemented with 10% (v/v) FCS. HCT116 and KMS-12-PE (DSMZ #ACC606) cells were cultured in RPMI1640 supplemented with 10% (v/v) FCS. OCI-AML3 cells were cultured in MEM alpha (GIBCO) supplemented with FBS 10% (v/v). and MV4;11 cells were cultured in RPMI (GIBCO) supplemented with FBS 10% (v/v). All media contained penicillin (100 IU/ml), streptomycin (100 _μ_g/ml), and L-glutamine (2 mM). Cells were cultured in humidified incubators maintained at 37 °C and 10% or 5% CO_2_. Cells were routinely screened for Mycoplasma contamination using MycoAlert Kit (Cat#LT07218, Lonza) as per manufacturer’s instructions. All cell lines were determined to be free of mycoplasma contamination. For transient transfection, constructs were introduced into cells by Lipofectamine 3000 transfection according to the manufacturer’s instructions.

### Generation of BAX knock-out AML cells using CRISPR-Cas9 gene editing

Stable Cas9-expressing human target cells were generated firstly through lentiviral transduction of target cells with FuCas9Cherry (Addgene Plasmid #70182), followed by cell sorting for mCherry expression using a FACSAria (Becton Dickinson: BD). Human stable Cas9 cell lines were subsequently transduced with lentiviral supernatants. Single guide (sg) RNAs targeting were synthesized and cloned into FgH1tUTG (Addgene Plasmid #70183), which permits doxycycline-inducible expression of the sgRNA and constitutive expression of a GFP reporter. All lentiviruses were produced in 293T cells and cells were transduced using established protocols. Stable Cas9-expressing target cells were generated firstly through transduction of target cells with FuCas9Cherry (Addgene Plasmid #70182), followed by cell sorting for mCherry expression using a FACSAria (Becton Dickinson: BD). Stable Cas9 cell lines were subsequently transduced with lentiviral supernatants. The loss of protein expression was confirmed by western blotting following culture of cells for 72 h in the presence of 5ug/ml doxycycline (Sigma Cat. No. 17086-28-1).

### Measurement of mitochondrial membrane potential

Protocol was modified based on publication(*27*). Cells were harvested and washed once with ice-cold PBS. Cells were re-suspended in 300 mM sucrose, 10 mM HEPES-KOH pH 7.7, 80 mM KCl, 1 mM EDTA, 1 mM EGTA, 5 mM Na succinate, 0.1% BSA with the concentration of 250000 per micro liter. Cells were treated with compounds for 10 min at room temperature, before the addition of BH3 peptide from human BIM and 0.001% digitonin. After 75 min incubation at room temperature, JC-1 dye was added to 100 nM. Samples were analyzed using a LSR-II flow cytometer (BD Biosciences).

### Colony formation assay

Cells were seeded in 6-well plate at a density of 1000 per well and cultured with indicated concentrations of treatments for 5 days. After treatment, cells were stained with 0.5% crystal violet (Cat#C0775, Sigma-Aldrich) for 20 min at room temperature and photographed using a ChemiDoc Imaging System (BioRad).

### Intra-cellular staining and flow cytometry

Following indicated treatments, cells were harvested and washed once with ice-cold PBS. For assessment of BAK and BAX activation or cytochrome *c* release, cells were treated with 20 µM caspase inhibitor QVD-OPh (Cat#OPH109, MP Biomedicals) for 30 mins prior to treatment with death stimuli. To assess BAK and BAX conformation change, cell pellets were fixed and then permeabilised using the eBioscience cell fixation and permeabilization kit (Cat#88882400, Thermo Fischer) according to the manufacturer’s instructions. Fixed cells were incubated with either conformation-specific BAK antibody (Clone#G317-2, 1:100, Cat# 556382, BD Pharmingen), BAX antibody (Clone#3, 1:100, Cat#610982, BD Pharmingen), followed by phycoerythrin (PE)-conjugated anti-mouse antibody (1:200, Cat#1031309, SouthernBiotech).

To measure cytochrome *c* release, cells were first permeabilized with 0.025% w/v digitonin in permeabilization buffer (20 mM HEPES pH 7.5, 100 mM sucrose, 2.5 mM MgCl_2_, 100 mM KCl) for 10 min on ice before fixation. Fixed cells were incubated with anti-cytochrome c-APC antibody (Clone#REA702, 1:50) (Cat#130111180, Miltenyi Biotec). Samples were analyzed using a LSR-II flow cytometer (BD Biosciences).

### Western blotting

Cells were permeabilized with 0.025% w/v digitonin for 10 min on ice and cytosol and heavy membrane fractions separated by centrifugation at 13000 *g* for 5 min at 4 °C. After treatments, membrane fractions were resuspended in ONYX lysis buffer (20 mM Tris-pH 7.4, 135 mM NaCl, 1.5 mM MgCl_2_, 1 mM EGTA, 10% (v/v) glycerol, supplemented with 1% (v/v) digitonin, 0.5 μg/ml complete protease inhibitor (Sigma-Aldrich)) for 30 min on ice. Following lysis, samples were centrifuged at 13,000 *g* at 4 °C for 10 min and supernatants were collected. Concentrations of cytosolic and heavy membrane proteins were determined using the Bradford protein assay and resolved by SDS-PAGE or BN-PAGE. Gels were transferred onto PVDF membrane and non-specific binding was blocked with 5% w/v non-fat milk in TBS-T (20 mM Tris-HCL pH 7.6, 137 mM NaCl, 0.1% Tween-20) for 1 h at room temperature. Membranes were incubated with primary antibody overnight at 4 °C. The membranes were washed with TBS-T three times, followed by incubating with appropriate HRP-conjugated secondary antibodies for 1 h in room temperature. The membranes were washed with TBS-Tween (0.01% v/v) prior to detection using enhanced chemiluminescent (ECL) reagent (Millipore) and imaged using a ChemiDoc Imaging System (BioRad). Protein levels were quantified by densitometric analysis (chemiluminescence) using the Image Lab 6.1 software.

### High-throughput screening

Assay-ready plates were prepared containing compound or controls. Compounds were screened at a final concentration of 20 μM, and all wells were backfilled with DMSO to final 0.5%. For the HTS, *BAK*^-/-^ KMS-12-PE cells were resuspended in RPMI supplemented with 2.5% FBS and seeded into 384-well white tissue culture plates (Greiner cat # 781098) at 20,000 cells per well in 50 μL using a Multidrop Combi reagent dispenser (ThermoScientific). The plates were incubated for 2 h (37°C, 5% CO_2_), before addition of 100 nL of ABT-737 (final concentration 250 nM, previously determined to reduce the viability of the *BAK*^-/-^ KMS-12-PE under these assay conditions to approximately 30%) by pintool transfer. Plates were incubated at 37°C, 5% CO_2_ for a further 22 h, then CellTiter-Glo^®^ (Promega) added to all wells to determine viability. Luminescence was read on an Envision plate reader (Perkin Elmer). Data were archived and analyzed in ActivityBase XE (IDBS) and TIBCO Spotfire Screening data quality was monitored by the Z’ for each assay plate. Plates were excluded from analysis if Z’<0.5 and the plates included had an average Z’ of 0.65. Data was normalised to percent viability relative to DMSO (100% viability) and EC_70_ ABT-737 (0% viability). Hits were selected on the criteria of % activity >3SD mean of samples in the same experiment.

### Chemical synthesis and characterizations

All reactions were performed under a dry nitrogen atmosphere. All used solvents were reagent grade and were provided by Sigma Aldrich. Analytical thin-layer chromatography was performed on Merck silica gel 60F254 aluminium-backed plates and was visualized by fluorescence under UV light. Chromatography was performed using a CombiFlash®Rf purification system (Teledyne, ISCO, Lincon, NE, USA), with pre-packed silica gel columns (particle size 0.040-0.063mm). The eluting systems used are indicated for each purification (see Supplemental Information).

### Advanced NMR Methods

The sample was prepared in a Micro NMR tube solved in 200 µL of Acetone-d_6_ (D, 99.9%) Cambridge Isotope. ^1^H and ^13^C NMR spectra data were acquired on a Bruker Ultrashield Plus 500MHz equipped with a with Prodigy BBO H&F Cryoprobe – 5 mm probe with frequency of 500.14 MHz for ^1^H and 12 µsec of pulse and frequency 100.62 MHz for ^13^C and 10 µsec of pulse. All the experiments and data were acquired and processed with Topspin Software 4.2.0 versions. Chemical shifts are reported in parts per million (ppm) on the δ scale and referenced to the appropriate solvent peak. The peak multiplicity is reported as follows: s (singlet), br s (Broad singlet), d (doublet), t (triplet), br t (broad triplet), q (quartet), qn (quintet), m (mutiplet), dd (doublet of doublet), dt (doublet of triplet), spet (septuplet).

#### Liquid chromatography mass spectroscopy (LCMS) was carried out using one of the following methods

Method A: Agilent G6120B MSD, using 1260 Infinity G1312B binary pump with 1260 detector Infinity G4212B DAD, the LC conditions as it follows, column; Poroshell 120 EC-C18, 2.1 x 30 mm 2.7 micron, injection volume: 1 µL; flow rate 1.0 mL/min; gradient 5-100 % of B over 3.8 min, (Solvent A: water, 0.1% formic acid; solvent B: ACN, 0.1% formic acid). MS conditions were as follows, ion source: single-quadrupole, ion mode: ES positive unless otherwise specified, source temperature: 150°C, desolvation temperature: 350°C, detection: ion counting, Capillary (KV)-3.00, Cone(V): 30, Extractor (V): 3, RF Lens (V): 0.1, Scan Range: 100-1000 Amu, Scan Time: 0.5 sec, Acquisition time: 4.1 min, Gas flow: desolvation L/hr-650, Cone L/hr-100.

Method B: Agilent G615B MSD, using 1260 Infinity II G7129B binary pump with 1290 detector Infinity G71172B DAD, the LC conditions as it follows, column Luna, Omega 3.0 µM PS C18 100 A, 50 x 2.1 mm; injection volume: 1 µL; flow rate 0.6 mL/min; gradient 5-100 %of B over 3.8 min, (Solvent A: water, 0.1% formic acid; solvent B: ACN, 0.1% formic acid). MS conditions were as follows, ion source: single-quadrupole, ion mode: ES positive unless otherwise specified, source temperature: 150°C, desolvation temperature: 350°C, detection: ion counting, Capillary (KV)-3.00, Cone(V): 30, Extractor (V): 3, RF Lens (V): 0.1, Scan Range: 100-1000 Amu, Scan Time: 0.5 sec, Acquisition time: 4.1 min, Gas flow: desolvation L/hr-650, Cone L/hr-100.

Method C: Waters AcQuity QDA mass detector and Waters 2998 photodiode array detector. For method C, HPLC conditions were as follows, column: XBridge TM prep C18 5 μm 19x100 mm; injection volume: 500 μL; flow rate 20 mL/min; gradient: 30-100% of B over 15 min, (Solvent A: water, 0.1% formic acid; solvent B: ACN, 0.1% formic acid). MS conditions were as follows, ion source: single-quadrupole, ion mode: ES positive unless otherwise specified, source temperature: 150°C, desolvation temperature: 350°C, detection: ion counting, Capillary (KV)-3.00, Cone(V): 30, Extractor (V): 3, RF Lens (V): 0.1, Scan Range: 150-1250 Amu, Scan Time: 0.5 sec, Acquisition time: 17 min, Gas flow: desolvation L/hr-650, Cone L/hr-100.

Method D: Waters 1525 Binary HPLC Pump and Waters 2998 photodiode array detector. Conditions: XBridge TM semi-prep C18 5 μm 19x100 mm column; injection volume: 300 μL; flow rate 15 mL/min; gradient: 400-100% of B over 28 min (Solvent A: water, 0.1% formic acid; solvent B: MeOH, 0.1% formic acid). UV detector using PDA detector. Acquisition time: 30 min.

HRMS analyses were carried out at the Analytical lab at the LaTrobe University Bundoora campus) on an Agilent 6530 TOF LC/MS Mass Spectrometer coupled to an Agilent 1290 Infinity (Agilent, Palo Alto, CA). All data were acquired, and reference mass corrected *via* a dual-pray electrospray ionisation (ESI) source. Acquisition was performed using the Agilent OpenLab ECM XT software. Compound **1** was purchased from Enamine (T0508-3203).

### Generation of stable cell lines

HEK293T cells were used as virus packaging cells. The viral constructs were first introduced into packaging cells by Lipofectamine 3000 transfection according to the manufacturer’s instructions. Viral supernatants were filtered and used to infect cells by spin-infection (2500 rpm centrifugation at 25 °C for 1 h) in the presence of 4 µg/mL polybrene (Cat#S2667, Sigma-Aldrich). Transduced cells were enriched either by antibiotic selection or expression of fluorescent protein using a BD FACSAria Fusion cell sorter.

### Cell viability assay

Cells were treated as indicated. After treatments, cells were harvested and re-suspended in KDS-BSS buffer with 2.5 to 5 µg/ml propidium iodide (PI) (Sigma Aldrich). Cell viability (PI negative) was assessed using an LSR-II flow cytometer (BD Biosciences).

For cell viability assay with AML cells, cell lines were plated at 2.5 × 10^5^ cells/mL (25000 cells per well) and treated with 6-point 10-fold serial dilution of compounds starting from 10 μM. Cell viability was determined after 48 h of treatment by FACS analysis of cellular exclusion of SYTOX Blue Dead Cell Stain (Life Technologies Cat No S34857) using an LSR-Fortessa (BD). All FACS data was analyzed using the FlowJo software.

### Recombinant protein expression and purification

Protocols for expression and purification of full-length human VDAC1 (UniProt: P21796) and VDAC2 (UniProt: P45880) were adapted from previous publications(*40*). Constructs with N-terminally His-tagged VDAC1 and VDAC2 were transformed into BL21 (DE3) and plated with 100 μg/mL ampicillin overnight at 37 °C. Single colonies were picked and inoculated into Luria Broth (LB) medium with 100 μg/mL ampicillin overnight at 37 °C with agitation. 15 mL of overnight starter culture was inoculated into Super Broth (SB) medium with 100 μg/mL ampicillin at 37 °C, until OD_600nM_ reached 1. Protein expression was induced with 1 mM IPTG. After another 3 h of culture, cells were harvested by centrifugation at 4500 rpm for 15 min and frozen at -80 °C for further experiments.

The bacterial cell pellet was lysed in lysis buffer (lysozyme, DNase I, 1% Triton X-100, 4 mM MgCl_2_ and 10 mM DTT) at room temperature for 20 min. Inclusion bodies were prepared after centrifugation at 10000 *g*, 15 min, 4 °C and washed 3 times with wash buffer I (50 mM Tris pH 8.0, 100 mM NaCl, 1 mM EDTA, 1 mM DTT, 0.5% Triton X-100 and protease inhibitors) and one time with wash buffer II (50 mM Tris pH 8.0, 1 mM EDTA, 1 mM DTT and protease inhibitors) in a Dounce homogenizer. Inclusion bodies were solubilized in 20 mM Tris pH 8.0, 6 M guanidinium, 0.5 mM EDTA and 1 mM DTT overnight at 4 °C. The insoluble pellet was removed by centrifugation at 20000 *g*, 30 min, 4 °C. Denatured proteins in supernatants were collected and frozen at -80 °C for further experiments.

Refolding of VDAC proteins was performed by dropwise dilution into refolding buffer (20 mM Tris pH 8.0, 300 mM NaCl, 2% N,N-Dimethyldodecylamine N-oxide (LDAO) and 1 mM TCEP) with stirring at 4°C. Protein aggregates were removed by centrifugation at 3500 *g*, 20 min, 4 °C.

Ni-NTA column was used for immobilized metal ion affinity chromatography (IMAC). Protein was loaded onto the Ni-NTA column followed by washes with more than 10 column volumes of 20 mM Tris pH 8.0, 300 mM NaCl, 0.1% LDAO, 1 mM TCEP at 4°C. Proteins were eluted with 20 mM Tris pH 8.0, 300 mM NaCl, 0.1% LDAO, 1 mM TCEP and 250 mM imidazole, pooled and concentrated.

Proteins were loaded onto a Superdex 200 Increase 10/300 GL size exclusion column (Cytiva) for further purification at 4°C. The column was pre-equilibrated with 20 mM Tris pH 8.0, 300 mM NaCl, 0.1% LDAO and 1 mM TCEP. After sample loading, proteins were eluted in the same buffer, and fractions were assessed on SDS-PAGE. Fractions with purified proteins were pooled, concentrated and stored at -80 °C.

### Drug affinity responsive target stability (DARTS) assay

Protocols for DARTS assay were adapted from previous publication(*31*). To perform DARTS assay on mitochondria, cells were first permeabilized with 0.025% w/v digitonin in permeabilization buffer (20 mM HEPES pH 7.5, 100 mM sucrose, 2.5 mM MgCl_2_, 100 mM KCl) for 10 min on ice. Membrane fractions were collected after centrifugation at 13000 *g* for 5 min at 4 °C. Membrane fractions containing mitochondria were re-suspended in permeabilization buffer without digitonin and incubated with compounds or DMSO at 30 °C for 30 min. Proteinase K was added in the sample, followed by incubation at 37 °C for 30 min. LDS loading dye with proteinase inhibitors were added to stop the reaction. Samples were heated at 95 °C for 8 min before loaded for SDS-PAGE followed by Western blotting.

Similar assay was performed on recombinant proteins. Purified proteins were retrieved from - 80 °C and thawed on ice. Proteins were incubated with compounds or DMSO in 20 mM Tris pH 8.0, 300 mM NaCl, 0.1% LDAO and 1 mM TCEP at 30 °C for 30 min. Proteinase K was added, followed by incubation at 37 °C for 30 min. LDS loading dye with proteinase inhibitors were added to stop the reaction. Samples were heated at 95 °C for 8 min before loaded for SDS-PAGE followed by Western blotting.

## Acknowledgments

Research is supported by grants and fellowships from the National Health and Medical Research Council, Australia (1043149, 1156024, DCSH; 2016894, RWB; 2009062, PEC; 1117089, GL; 2018071, AHW; 2004446, GD), Bodhi Education Fund (GD), Victorian Cancer Agency MCRF Fellowship (19011, DMM). The Australian Drug Discovery Library (ADDL) was funded by MTPConnect which is supported by the Australian Government, Department of Industry, Science and Resources. We acknowledge Compounds Australia (www.compoundsaustralia.com) for their provision of specialized compound management and logistics research services to the project. This work was supported by operational infrastructure grants through the Australian Government Independent Research Institute Infrastructure Support Scheme (9000587) and the Victorian State Government Operational Infrastructure Support, Australia.

## Funding

National Health and Medical Research Council, Australia 1043149 (DCSH)

National Health and Medical Research Council, Australia 1156024 (DCSH)

National Health and Medical Research Council, Australia 2016894 (RWB)

National Health and Medical Research Council, Australia 2009062 (PEC)

National Health and Medical Research Council, Australia 1117089 (GL)

National Health and Medical Research Council, Australia 2018071 (AHW)

National Health and Medical Research Council, Australia 2004446 (GD)

Bodhi Education Fund (GD)

Victorian Cancer Agency MCRF Fellowship 19011 (DMM)

## Author contributions

Conceptualization: KL, MvD, GD, GL, BTK, PEC, AHW, DCSH

Methodology: KL, MvD, DMM, YK, KLo

Investigation: YQY, FS, AG, MJ, TEL, MDS, MXL, JG, ZY, RWB, DMM, YK, KLo

Supervision: GD, MvD, GL

Writing—original draft: KL, DMM, YK, KLo, GD, MvD, GL

Writing—review & editing: PEC, DCSH, GD, MvD, GL, KL, BTK

## Competing interests

GD, PEC, RB, DCSH, MvD, GL, YQY, DMM, FS, YK, AG, AWR, RWB, PEC, KLo and AHW are employees of the Walter and Eliza Hall Institute of Medical Research that receives milestone payments related to sales of venetoclax. AHW provides medical advice and receives honoraria and research funding to the institution from AbbVie.

## Data and materials availability

All data are available in the main text or the supplementary materials.

## Supplementary Figure Legends

**Fig. S1. High-throughput cell-based phenotypic screening and discovery of WEHI-3773. (A)** Western blot analysis of whole cell lysates of KMS-12-PE cells in this study. Representative blots of at least 3 independent experiments are shown. **(B)** Dose–response curves for ABT-737 on KMS-12-PE cells. Cells were treated with different concentrations of ABT-737. 48 h later, cells were collected, and cell death was quantified by PI uptake and flow cytometry. Data are presented as mean and S.E.M. of 3 independent experiments. **(C)** Dose– response curves for Compound 1 on *BAX*^-/-^ KMS-12-PE quantified with CellTiter-Glo^®^ 2.0. Data are presented as mean and S.E.M. of 2 independent experiments. **(D)** Dose–response curves for Compound **1** on WT KMS-12-PE quantified using CellTiter-Glo^®^ 2.0. Data are presented as mean and S.E.M. of 2 independent experiments. **(E and F)** Decomposition profile of Compound **1** (**E**) and Compound **2** in methanol and dichloromethane (**F**). **(G)** Synthesis of WEHI-3773 starting from Compound **2**.

**Fig. S2. Characterization of HeLa cell lines and colony formation with HCT-116 cells. (A)** Western blot analysis of whole cell lysates of Hela used in this study. Representative blots of at least 3 independent experiments are shown. **(B)** Dose–response curves for ABT-737 plus S63845 on HeLa cells. Cells were treated with different concentrations of ABT-737 plus a fixed concentration of S63845 (500 nM). 24 h later, cells were collected, and cell death was quantified by PI uptake and flow cytometry. Data are presented as mean and S.E.M. of 3 independent experiments. **(C)** Representative images of HCT-116 colonies with indicated treatments for 14 days. **(D)** Numbers of colonies quantified. Data are presented as mean and S.E.M of 3 independent experiments.

**Fig. S3. WEHI-3773 alone does not activate BAK or BAX S184L in the absence of apoptotic stressors.** Dose response viability curves of KMS-12-PE (**A**) and HeLa (**B**) measured by PI uptake and flow cytometry. Cells were treated with WEHI-3773 for 48 h before measurement. Data is mean ± S.E.M. of 3 independent experiments. **(C)** Western blot analysis of *BAX^-/-^* HeLa treated with WEHI-3773 or ABT-737 plus S63845 at indicated concentrations for 2 h, followed by blue native PAGE. Representative blots of at least 3 independent experiments are shown. **(D)** Intra-cellular staining and flow cytometry to measure conformationally activated BAK within *BAX*^-/-^ HeLa. **(E)** Dose response viability curves of MEF measured by PI uptake and flow cytometry. Cells were treated with WEHI-3773 for 24 h before measurement. Data is mean ± S.E.M. of 3 independent experiments.

**Fig. S4. Target identification in cells. (A)** Dose response viability curves of KMS-12-PE cells measured by PI uptake and flow cytometry. Cells were pre-treated with WEHI-9625 for 2 h first, then EC_70_ or EC_50_ concentrations of ABT-737 was added for another 46 h before measurement. Data are mean ± S.E.M. of 3 independent experiments. **(B)** Western blot analysis of mitochondria treated with DMSO or WEHI-3773 (400 nM) for 30 min, followed by addition of increasing concentration of proteinase K. Representative blots of 3 independent experiments are shown. **(A) (C)** Immunoblots of (**B**) were quantified and normalized to the sample untreated with proteinase K. Data are mean ± S.E.M. of 3 independent experiments.

**Fig. S5. Characterization of VDAC2 as a target for WEHI-3773. (A)** Purification of VDAC1. Marked fractions from size exclusion chromatography were analyzed by SDS-PAGE and stained by Coomassie blue. Fractions b to e were collected, pooled, concentrated and flash frozen for further experiments. **(B)** Purification of VDAC2. Fractions *a* to *e* were collected, pooled, concentrated and flash frozen for further experiments. **(B)** The sequence alignment was created using the web services ESPript 3.0. Identical and similar residues are boxed in red and white, respectively.

**Fig. S6. Effects of WEHI-3773 on resistant OCI-AML3. (A)** Dose–response curves for OCI-AML3 cells resistant to indicated BH3 mimetics. OCI-AML3 DMSO cells served as the control. Cells were treated with the indicated concentration of BH3 mimetics prior to assessment of cell viability after 48 h. Data is mean ± S.D. of 3 independent experiments. **(B)** OCI-AML3 cells resistant to S63845 (MCL-1i R) were treated with the indicated concentration of BH3 mimetics with or without WEHI-3773 (200 nM). Cell viability was determined after 48 h by flow cytometric enumeration of sytox blue exclusion. Collated data is mean ± S.D. of 3 independent experiments. **(C)** OCI-AML3 cells resistant to venetoclax plus S63845 (BCL2i + MCL1i R) were treated with the indicated concentration of BH3 mimetics with or without WEHI-3773 (200 nM). Cell viability was determined after 48 h by flow cytometric enumeration of Sytox blue exclusion. Collated data is mean ± S.D. of 3 independent experiments.

