## Supplementary Figures for "Differential regulation of BAX and BAK apoptotic activity revealed by a novel small molecule"

Figure S1

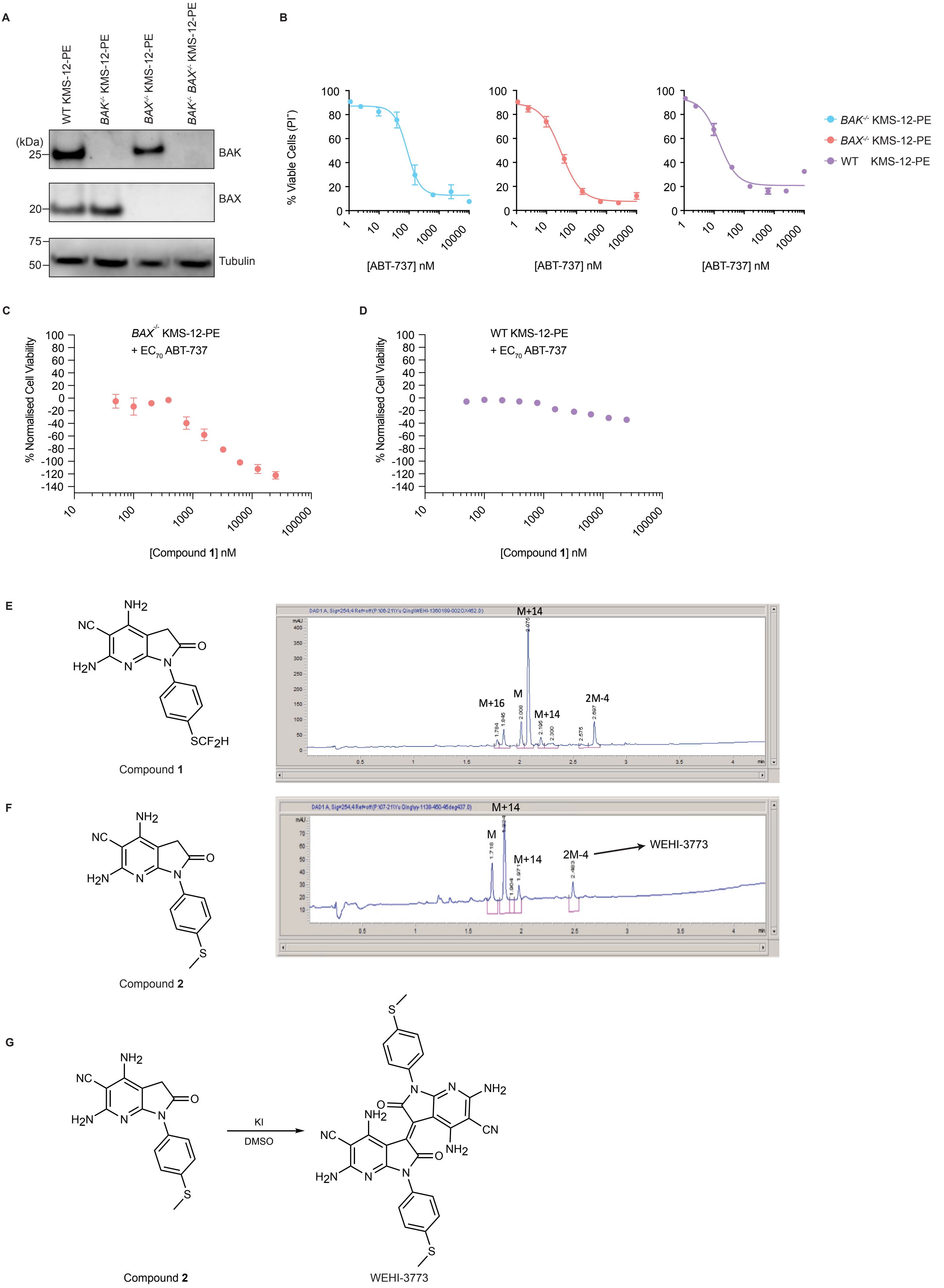
A
WT KMS-12-PE
BAK-/- KMS-12-PE
BAX-/- KMS-12-PE
BAK-/- BAX-/- KMS-12-PE
(kDa)
25
20
75
50
BAK
BAX
Tubulin
B
% Viable Cells (PI-)
[ABT-737] nM
BAK-/- KMS-12-PE
BAX-/- KMS-12-PE
WT KMS-12-PE
C
BAX-/- KMS-12-PE
+ EC70 ABT-737
% Normalised Cell Viability
[Compound 1] nM
D
WT KMS-12-PE
+ EC70 ABT-737
% Normalised Cell Viability
[Compound 1] nM
E
Compound 1
M+14
M+16
M
M+14
2M-4
F
Compound 2
M+14
M
M+14
2M-4
WEHI-3773
G
KI
DMSO
Compound 2
WEHI-3773

Figure S2

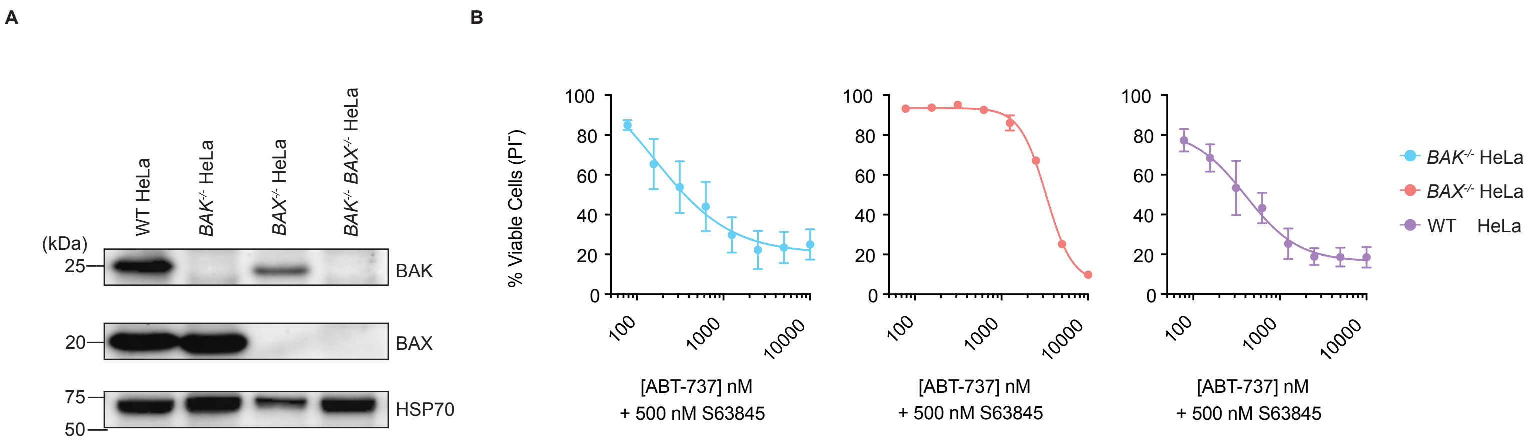
A
WT HeLa
BAK-/- HeLa
BAX-/- HeLa
BAK-/- BAX-/- HeLa
(kDa)
25
20
75
50
BAK
BAX
HSP70
B
% Viable Cells (PI-)
100
80
60
40
20
0
100
1000
10000
[ABT-737] nM
+ 500 nM S63845
BAK-/- HeLa
BAX-/- HeLa
WT HeLa

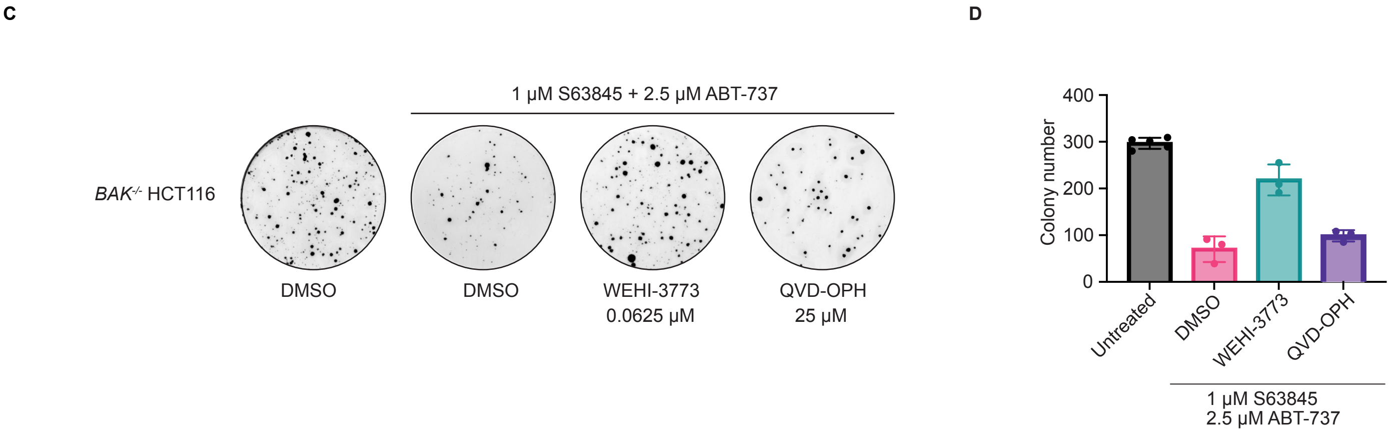
C
1 µM S63845 + 2.5 µM ABT-737
BAK-/- HCT116
DMSO
DMSO
WEHI-3773
0.0625 µM
QVD-OPH
25 µM
D
Colony number
400
300
200
100
0
Untreated
DMSO
WEHI-3773
QVD-OPH
1 µM S63845
2.5 µM ABT-737

Figure S3

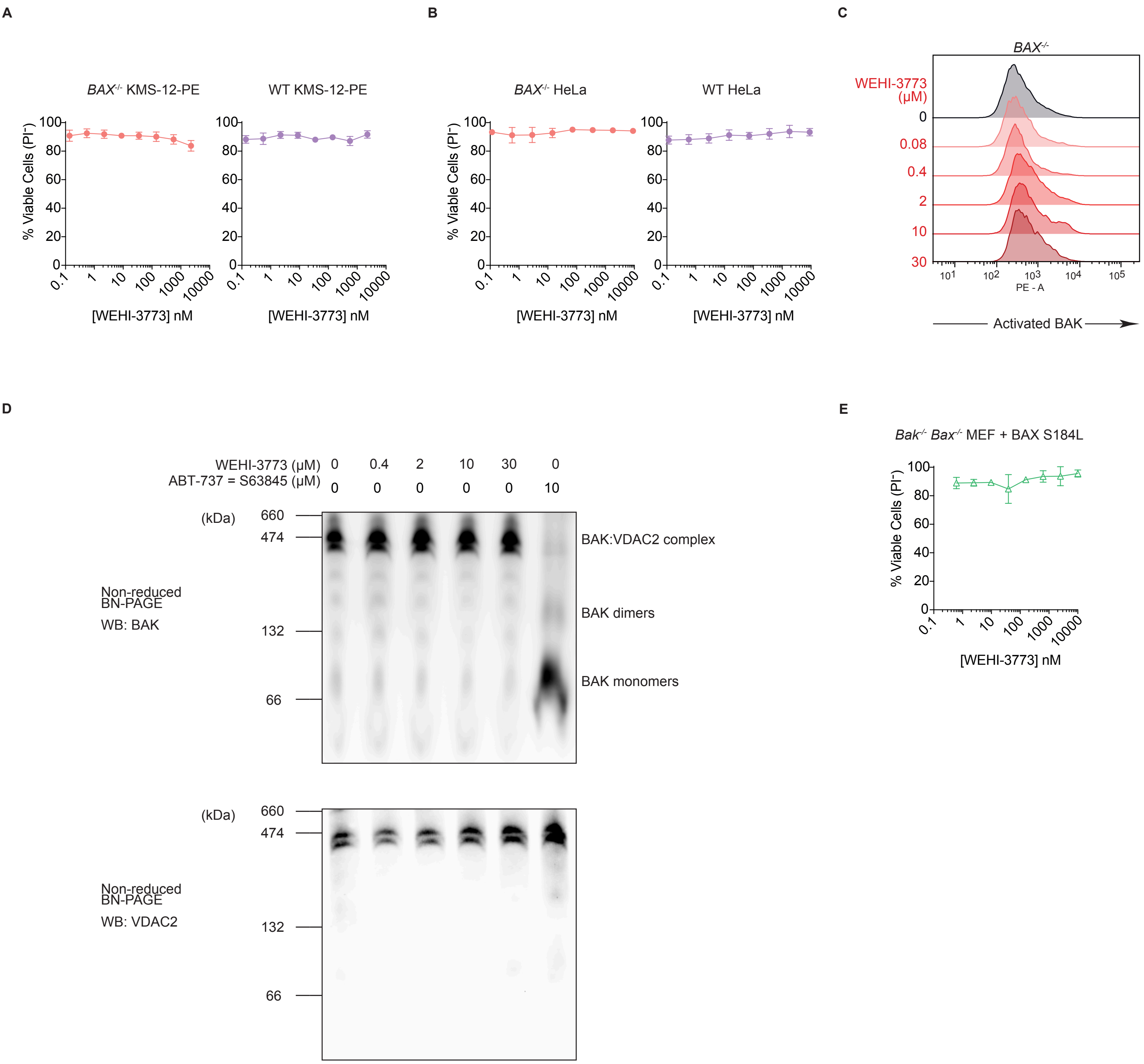
A
BAX-/- KMS-12-PE
WT KMS-12-PE
% Viable Cells (PI-)
[WEHI-3773] nM
B
BAX-/- HeLa
WT HeLa
% Viable Cells (PI-)
[WEHI-3773] nM
C
BAX-/-
WEHI-3773 (μM)
0
0.08
0.4
2
10
30
PE - A
Activated BAK
D
WEHI-3773 (μM) 0 0.4 2 10 30 0
ABT-737 = S63845 (μM) 0 0 0 0 0 10
(kDa)
660
474
132
66
Non-reduced BN-PAGE
WB: BAK
BAK:VDAC2 complex
BAK dimers
BAK monomers
Non-reduced BN-PAGE
WB: VDAC2
E
Bak-/- Bax-/- MEF + BAX S184L
% Viable Cells (PI-)
[WEHI-3773] nM

Figure S4

A

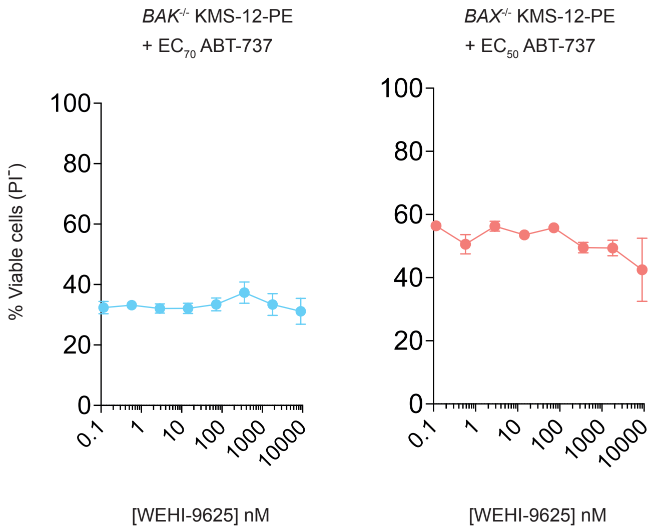


B

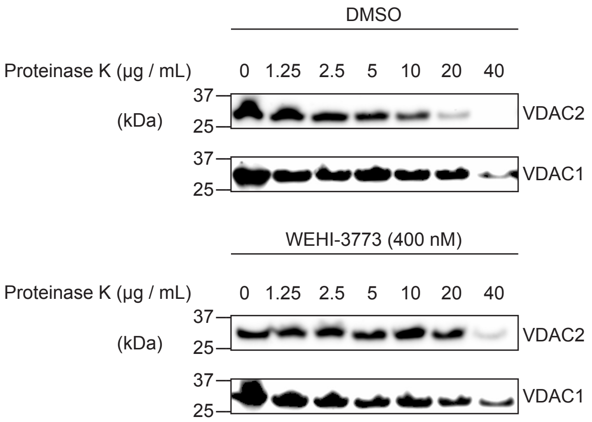


C

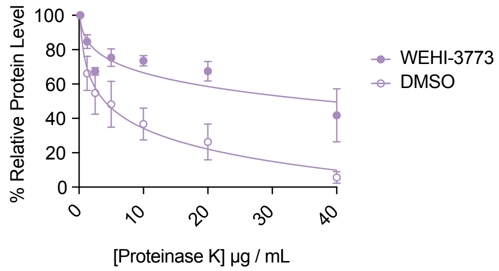

Figure S5

A

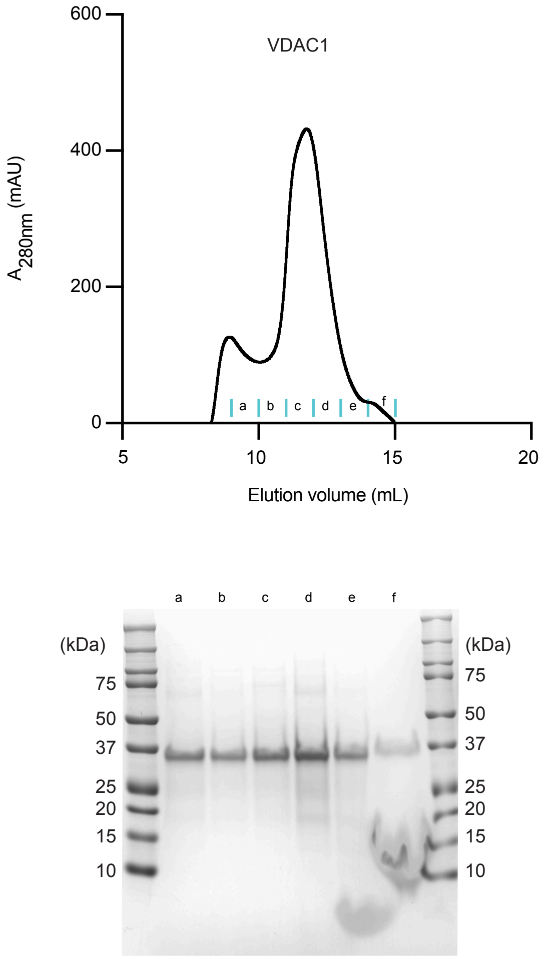


B

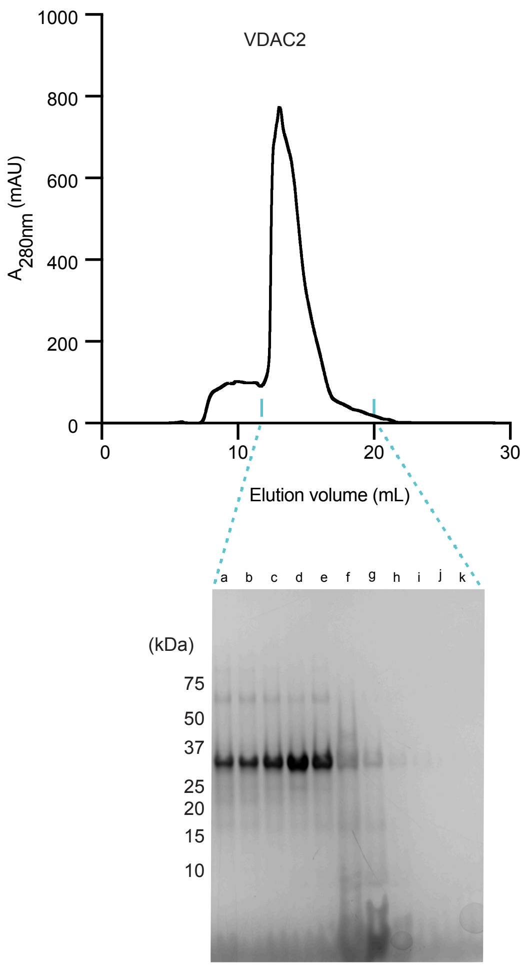


C

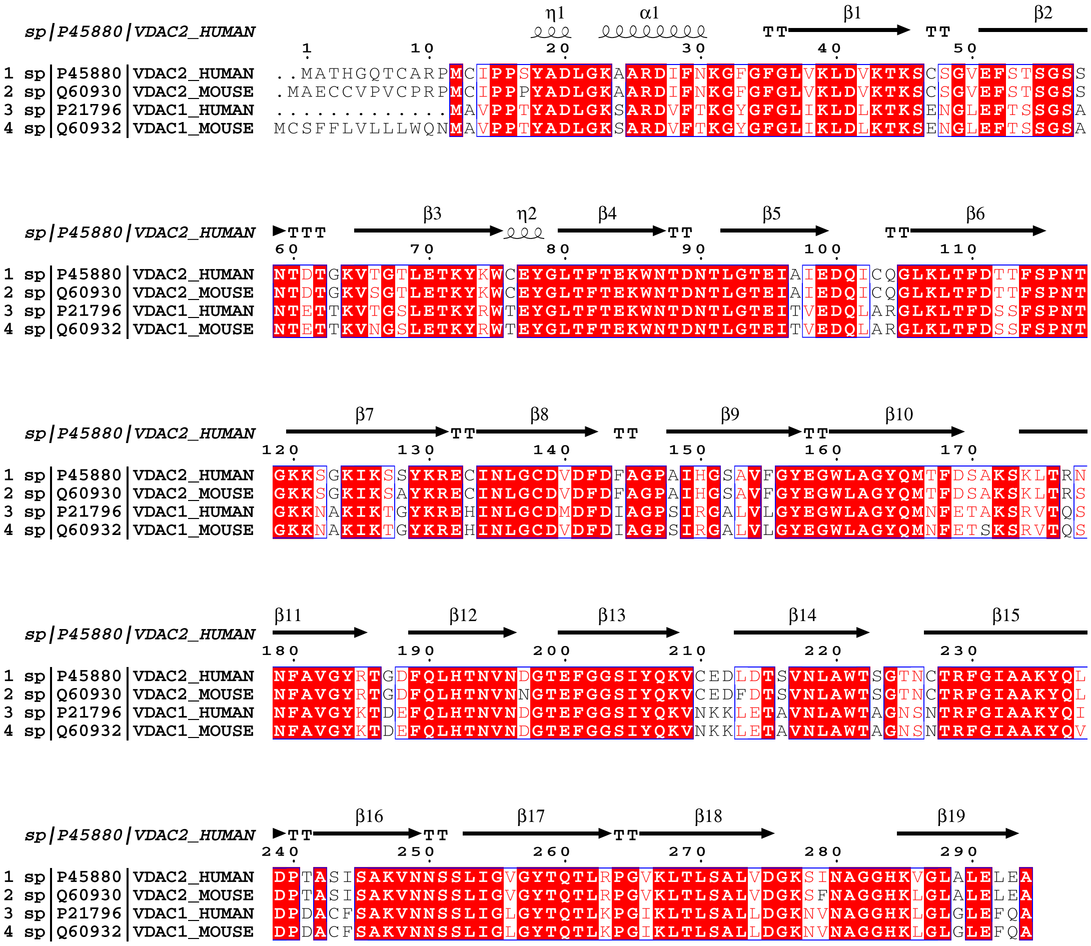

Figure S6

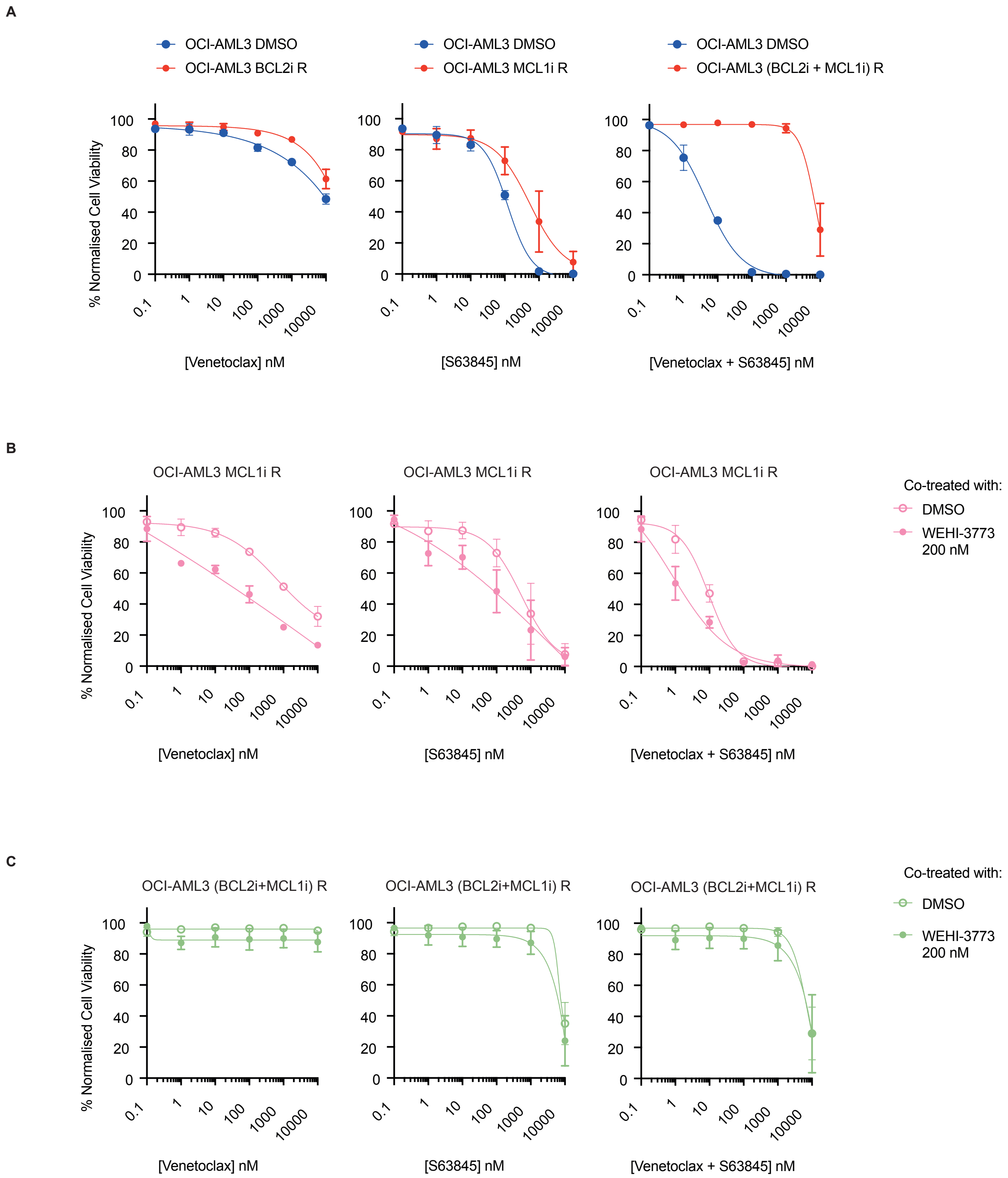
A
OCI-AML3 DMSO
OCI-AML3 BCL2i R
OCI-AML3 DMSO
OCI-AML3 MCL1i R
OCI-AML3 DMSO
OCI-AML3 (BCL2i + MCL1i) R
% Normalised Cell Viability
[Venetoclax] nM
[S63845] nM
[Venetoclax + S63845] nM
B
OCI-AML3 MCL1i R
OCI-AML3 MCL1i R
OCI-AML3 MCL1i R
Co-treated with:
DMSO
WEHI-3773 200 nM
% Normalised Cell Viability
[Venetoclax] nM
[S63845] nM
[Venetoclax + S63845] nM
C
OCI-AML3 (BCL2i+MCL1i) R
OCI-AML3 (BCL2i+MCL1i) R
OCI-AML3 (BCL2i+MCL1i) R
Co-treated with:
DMSO
WEHI-3773 200 nM
% Normalised Cell Viability
[Venetoclax] nM
[S63845] nM
[Venetoclax + S63845] nM

**Chemical synthesis and characterization**

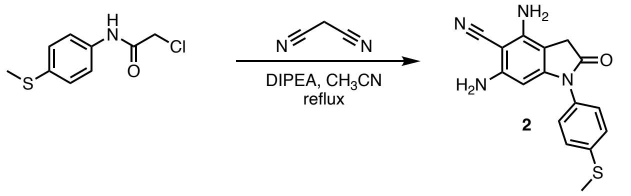


*Synthesis of 4,6-diamino-1-(4-methylsulfanylphenyl)-2-oxo-3H-pyrrolo[2,3-b]pyridine-5-carbonitrile* (**2**). In a round bottom flask, propanedinitrile (1,1 g, 16 mmol) was dissolved in acetonitrile (10 mL). Di-isopropylethylamine (4.54 mL, 26.1 mmol) was then added, and the reaction was heated to 50 ºC and stirred for 30 minutes. To this solution was slowly added 2-chloro-N-(4-methylsulfanylphenyl)acetamide (1,4 g, 6.51 mmol) dissolved in acetonitrile (15 mL). The reaction mixture was warmed to reflux and stirred for 5 hours. After that time, the reaction was cooled to room temperature at which point a precipitate formed. The solid was filtered and dried to give 4,6-diamino-1-(4-methylsulfanylphenyl)-2-oxo-3H-pyrrolo[2,3-b]pyridine-5-carbonitrile **2** (446 mg, 22% yield) . $^{1}$H NMR (300.13 MHz, $CDCl_3$) δ 7.43-7.39 (d, $J$ = 12.0 Hz, 4H), 7.32-7.28 (d, $J$ = 12.0 Hz, 4H), 6.02 (s, 4H, $NH_2$), 5.91 (s, 4H, $NH_2$), 2.81 (s, 6H). LCMS (ES+), m/z = 312.0 $[M+H]^+$.

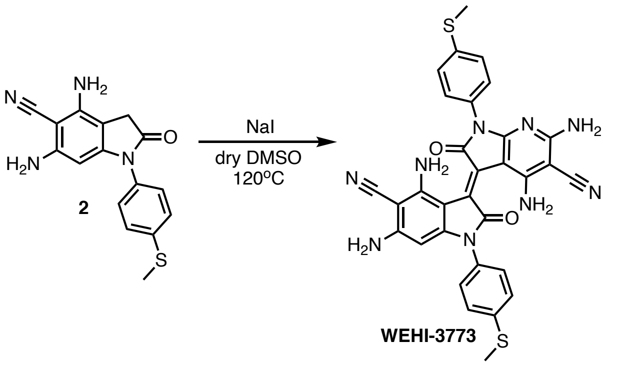


*Synthesis of (3E)-4,6-diamino-3-[4,6-diamino-5-cyano-1-(4-methylsulfanylphenyl)-2-oxo-pyrrolo[2,3-b]pyridin-3-ylidene]-1-(4-methylsulfanylphenyl)-2-oxo-pyrrolo[2,3-b]pyridine-5-carbonitrile* (**WEHI-3773**). In an oven dry round bottom flask was added 4,6-diamino-1-(4-methylsulfanylphenyl)-2-oxo-3H-pyrrolo[2,3-b]pyridine-5-carbonitrile **2** (100 mg, 0.32 mmol) and sodium iodide (241 mg, 1.6 mmol). The round bottom flask was evacuated and refilled with nitrogen three times. Dry DMSO (5 mL) was added to the round bottom flask and was fitted with a drying tube. The reaction was then heated to 120°C for 3 hours. The reaction mixture was cooled to room temperature and water (10 mL) was added to the round bottom flask, a dark precipitate was formed and filtered. The dark brown solid was dried in a vacuum oven overnight and purified using column chromatography, using 5 % MeOH/DCM. The fractions containing the expected compound were analyzed and combined. Further purification was performed using preparative HPLC Method C followed by Method D, to give (3E)-4,6-diamino-3-[4,6-diamino-5-cyano-1-(4-methylsulfanylphenyl)-2-oxo-pyrrolo[2,3-b]pyridin-3-ylidene]-1-(4-methylsulfanylphenyl)-2-oxo-pyrrolo[2,3-b]pyridine-5-carbonitrile **WEHI-3773** (5.0 mg, 2.5% yield) as a dark purple solid. $^{1}$H NMR (500.14 MHz, Acetone-$d_6$) δ 7.51-7.49 (d, *J* = 10.0 Hz, 4H), 7.41-7.39 (d, *J* = 10.0 Hz, 4H), 6.60 (s, 4H, $NH_2$), 6.49 (s, 4H, $NH_2$), 2.57 (s, 6H).$^{13}$C NMR (100.62 MHz, Acetone-$d_6$) δ 170.73, 163.86, 162.47, 155.35, 140.31, 132.77, 130.33, 128.04, 121.53, 118.09, 97.00, 72.46, 16.59. LCMS (ES+), m/z = 618.14 $[M+H]^+$. HRMS (+ESI), m/z = 619.1453$[M+H]^+$ (calculated: 619.1448).

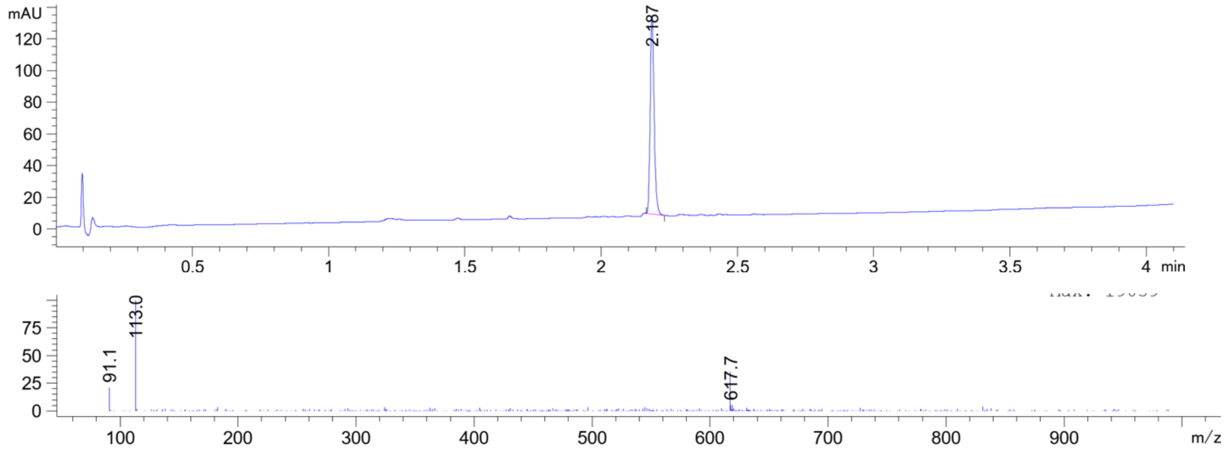

**WEHI-3773 – $^{1}$H NMR**

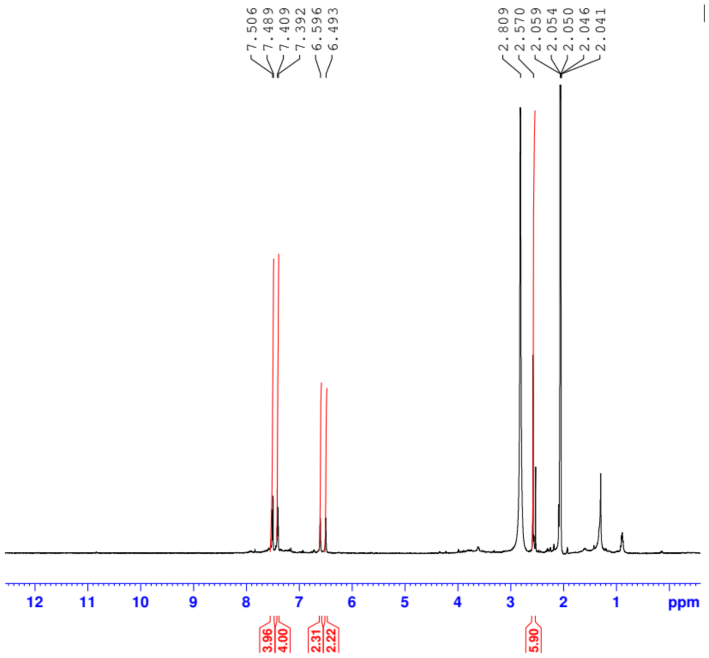


**WEHI-3773 – $^{13}$C NMR**

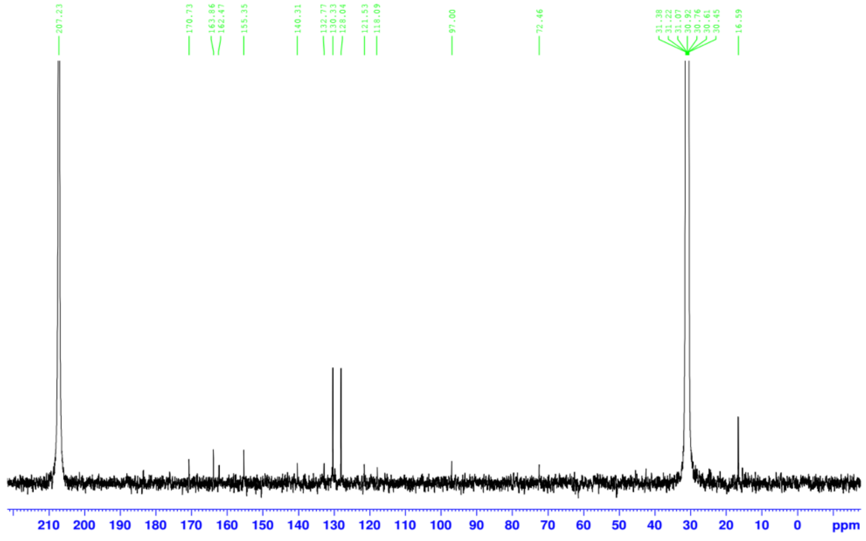

**HBMC**

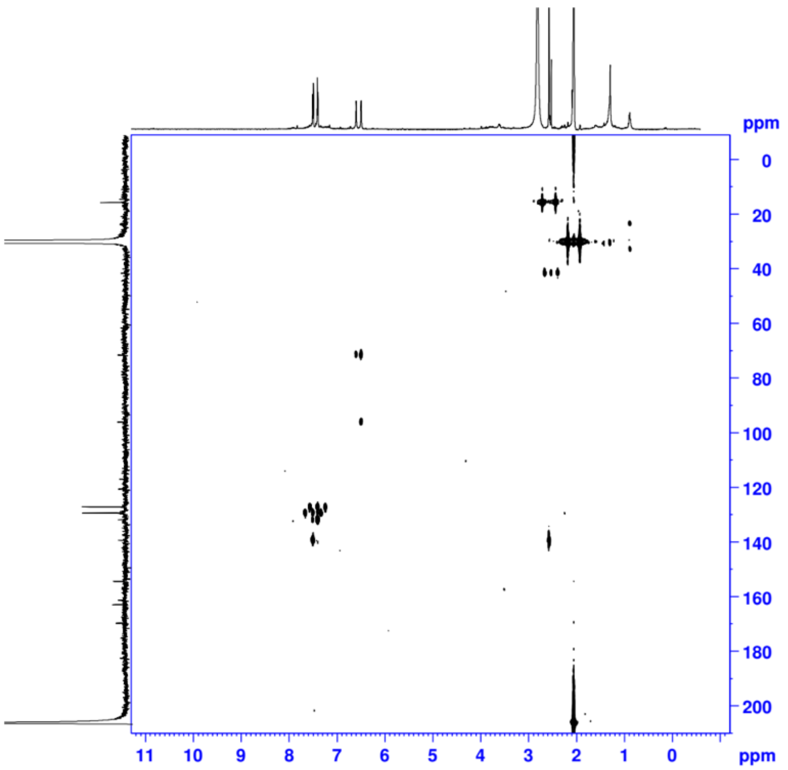


**HSQC**

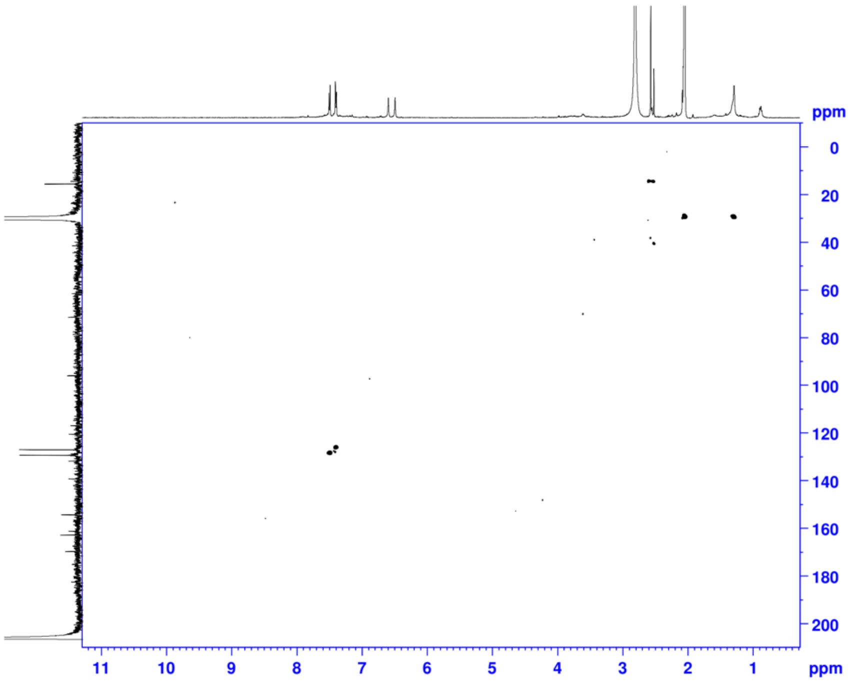


From 1D and 2D spectral data, the $^{13}C$ assignment is shown in the table below.

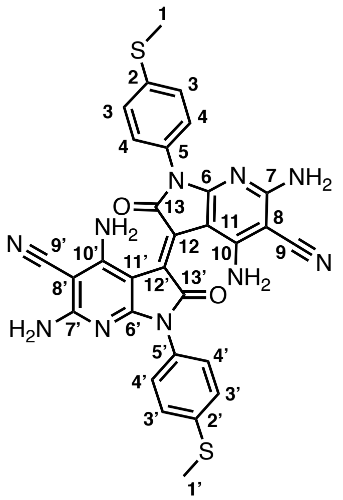


| Carbon | δ (ppm) |
|---|---|
| 1/1' | 16.59 |
| 2/2' | 140.31 |
| 3/3' | 130.33 |
| 4/4' | 128.04 |
| 5/5' | 132.77 |
| 6/6' | 163.86 |
| 7/7' | 162.47 |
| 8/8' | 72.46 |
| 9/9' | 118.09 |
| 10/10' | 155.35 |
| 11/11' | 97.0 |
| 12/12' | 121.53 |
| 13/13' | 170.73 |
